# Multi-stage single-cell atlas of *Botryllus schlosseri* asexual development unveils dedifferentiating bud founder cells

**DOI:** 10.1101/2025.05.27.655565

**Authors:** Marie Lebel, Tiphaine Sancerini, Sharon Rabiteau, Solène Marchal, Pragati Sharma, Tal D. Scully, Estelle Balissat, Laurel S. Hiebert, Anthony W. De Tomaso, Allon M. Klein, Alexandre Alié, Stefano Tiozzo

## Abstract

Colonial tunicates are the only chordates capable of forming fully functional bodies from somatic tissues through non-embryonic development, known as budding. In *Botryllus schlosseri,* this agametic process, termed peribranchial budding, generates new zooids in a stereotyped, cyclical manner from a cluster of cells within the peribranchial epithelium. Despite detailed morphological characterization, the molecular and cellular underpinnings of budding initiation remain poorly understood. Here, we present the first single-cell transcriptomic atlas of *B. schlosseri* peribranchial budding, encompassing multiple stages from pre-budding to near-mature zooid. This high-resolution atlas captures the diversity of cell types involved in budding and their dynamics enabling precise cluster annotation and lineage trajectory inference. While circulating mesenchymal cells exhibit transcriptional hallmarks of stem-like states, we found no definitive evidence of broad contribution to bud onset and early morphogenesis beyond hematopoietic and gonadal lineages. Instead, we identified a distinct founder cell population arising from peribranchial epithelium, marked by a unique transcriptional profile and progressive acquisition of developmental potency. This supports a model in which budding is initiated by dedifferentiation of committed epithelial cells rather than activation of multi- or pluripotent stem cells. Furthermore, we highlight signaling pathways, including GNRHR-like receptors, that may couple metabolic state with developmental progression. Altogether, our data suggest that peribranchial budding in *B. schlosseri* is driven by potential reprogramming within epithelial tissues. This work provides a foundational resource for studying non-embryonic development and the evolution of regenerative strategies in chordates.

## Introduction

Colonial tunicates are the only chordates capable of developing a fully functional body from somatic cells, either through cyclical agametic propagation or via whole-body regeneration in response to extensive injury. These forms of non-embryonic development are collectively referred to as budding (1, 2). When integrated into the agametic life-cycle, budding typically results in newly formed individuals that remain physically connected to the parent zooid, ultimately giving rise to colonies composed of up to hundreds of genetically identical individuals.

Tunicate phylogeny suggests that budding is an evolutionarily labile trait, having evolved independently multiple times (3, 4). In contrast to the highly conserved embryonic development observed across major tunicate orders, budding displays remarkable diversity: it can originate from a variety of cell types and tissues, often nonhomologous, proceeds through distinct developmental trajectories, and exhibits variable interactions between epithelial and mesenchymal cells, even within the same species (5). While the morphological diversity of tunicate budding has been recorded in scattered literature for over two centuries of (reviewed in (4)), much of the current understanding stems from more recent investigations in a few model species, particularly within the Styelidae family (5). Among these, *Botryllus schlosseri* has emerged as leading laboratory model over the past decades, not only for the study of budding (6, 7), but also for investigating biological processes such as allorecognition and chimerism (8, 9). In *B. schlosseri*, agametic propagation occurs via a highly stereotyped mode of budding known as palleal or peribranchial budding (1, 7). During this process, buds arise from specific regions of the peribranchial epithelium of the parental zooid and progress through stereotyped morphogenetic stages (Fig. 1A; see Methods; (1, 10)). At a given time, a colony harbors three synchronized asexual generations: the adult filtering zooids, their primary buds, and the secondary buds (or budlets) arising from the primary buds. The colonies periodically undergo a synchronized apoptotic clearance of the adults (take-over) that are replaced by their mature primary buds (stage D, Fig. 1A; (1)). Peribranchial budding is a continuous, lifelong process that, unlike sexual reproduction, bypasses embryonic stages and metamorphosis, and enables the colony to grow and propagate.

**Figure 1.**
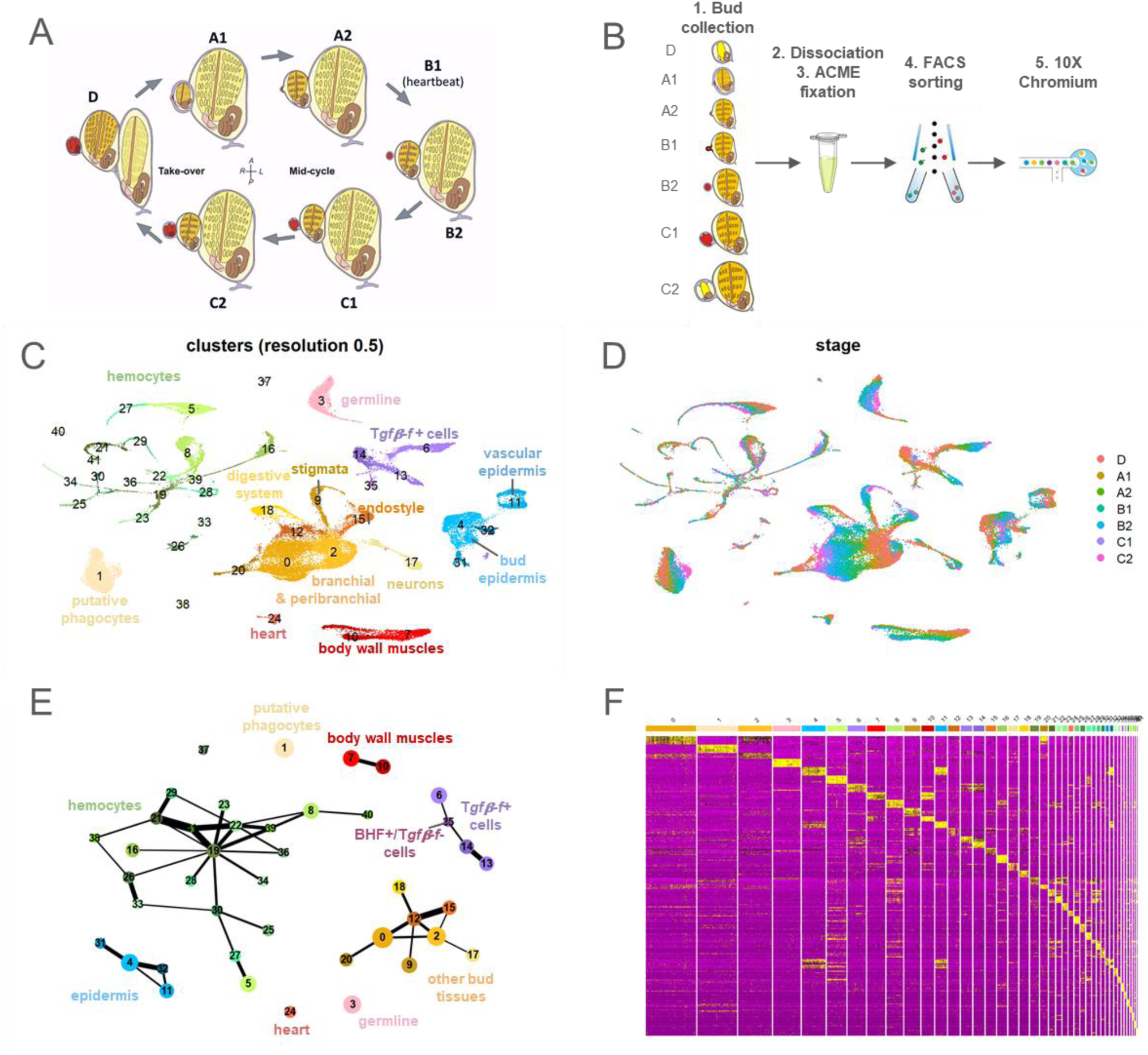
A single cell atlas over the development of the bud. **(A)** Schematic representation of *B. schlosseri* peribranchial budding cycle, with staging according to (8) (modified from Manni et al., 2014 (1)). **(B)** Experimental workflow (*B. schlosseri* schematics modified from Manni et al., 2014 (1)). **(C)** UMAP projection of the full budding dataset showing the 42 clusters at resolution 0.5. **(D)** Same UMAP as in (C), with cells labeled by stage. **(E)** Abstracted graph (PAGA) of the full dataset clusters (PAGA threshold 0.155), showing 9 major groups. **(F)** Heatmap of the top ten differentially expressed genes per cluster.

The anatomy and morphological stages of budding in *Botryllus schlosseri* have been well described (1), and several studies have investigated the molecular mechanisms underlying regulation of early cell commitment (11–13), establishment of developmental axes and symmetry (14–16), gametogenesis (17, 18), angiogenesis (19–21) and the differentiation of specific tissues such as muscle and nervous lineages (22, 23). Parallel to these findings, various hypotheses have emerged regarding the identity and nature of the cells that initiate the budding process. One long-standing hypothesis posits that the development of asexually derived bodies originates from a pool of stem cells functionally analogous to embryonic stem cells (24). This is supported by studies showing the engrafting and long-term contribution of circulating cells to the gonads or some somatic tissues (25). Similarly, Voskoboynik and colleagues used *in vivo* cell labeling, engraftment, and time-lapse imaging, to identify cells with stem-like properties in a not clearly defined region antero-ventral to the endostyle (26). These cells exhibit proliferation and migratory behaviors, and seem to contribute to the generation of specific organs in developing buds, yet they do not participate in germline formation. The same group also identified discrete structures named ‘cell islands’ proposed to serve as a potential niche housing *Piwi*-positive cells proposed to contribute to both germline and somatic stem cell populations during budding (27). Studies in closely related species, such as *Botrylloides violaceus* and *Botrylloides diegensis*, have similarly reported the presence of circulating undifferentiated cells involved in injury-triggered whole-body regeneration, a phenomenon referred to as vascular budding (28, 29). An alternative hypothesis proposes that in colonial tunicates, the remarkable flexibility of tissue remodeling observed in whole-body regeneration and agametic reproduction is driven by two complementary systems: the dedifferentiation of epithelial cells and the epithelial transformation of circulating cells, bypassing the need for activation of multi- or pluripotent stem cells (reviewed in (30)).

Despite extensive descriptions of morphology, anatomy, and ontogeny of bud development in *B. schlosseri* (1, 31), and mounting insights into early tissue specification and patterning with the identification of putative stem-cell niches, the identity of the founder cells and the molecular mechanisms that initiate peribranchial budding, particularly the acquisition of developmental competence in already committed tissues, remain largely elusive.

In this study, we present the first multi-stage single-cell transcriptomic atlas of tunicate agametic development, capturing the successive phases of peribranchial budding in *Botryllus schlosseri*, from the earliest stage prior to budlet formation until the emergence of a nearly mature zooid. This dataset enabled us to pinpoint the epithelial origin of budding cells by identifying and characterizing the transcriptomic profile of the founder population in the peribranchial epithelium. Moreover, the high-resolution atlas provides a functional resource for exploring the cellular and molecular mechanisms underlying non-embryonic development in chordates. By offering a detailed portrait of the diverse cell types engaged in budding, along with robust gene expression profiles that facilitate accurate cluster annotation, we open the door to revisiting long-standing developmental hypotheses within a comprehensive transcriptomic and genomic framework.

## Material and Methods

### Animal collection and husbandry

To ensure genetic homogeneity for the budding samples, young oozoids were collected on glass slides from the port of Villefranche-sur-Mer (43.697882, 7.30735), then isolated in small containers (∼1 L) at 18°C to let them grow for a few weeks. They were fed with a mix of live *Tisochrysis lutea*, live *Chaetoceros gracilis*, ISO800 and Shellfish Diet 1800 (Reed Mariculture). The water changed about once a day. Once the colonies had 10-20 blastozooids, they were transferred to a cold marine culture system (12°C) described previously (32). One of these isogenic colonies (labelled as FC*) was chosen to perform all the dissociations. For the circulating cell samples, animals were collected in Santa Barbara, California, USA. They were kept in a semi flow through mariculture system with 0.5µm-filtered seawater at 16-18 °C, and they were fed twice a day with cultures of the following live microalgae: *Dunaliella sp.* (Carolina Biological CAT#152160), *Isochrysis sp.* (Carolina Biological CAT#153180), *Nannochloropsis sp.* (Carolina Biological CAT#153220), *Tetraselmis sp.* (Carolina Biological CAT#152610). Colonies were attached to glass slides with sewing thread and allowed to grow onto slides. The slides were kept vertically in aquarium tanks. Slides were cleaned every few days to a week with a paint brush, toothpick, and razor blade as needed to remove decay, debris, and microbe/parasite growth. Colonial animals were allowed to grow for a couple of months in the lab. They were then shipped overnight to Boston, at which point they were kept in artificial sea water (ASW, Instant Ocean SS15-10) at 17 °C.

### Sample collection and dissociation

Animals were staged according to (10) under a stereomicroscope. Each budding cycle begins with the formation of a new body. A new budlet initially appears as a disc of thickened cells (stage A1) in a specific region of the primary bud’s peribranchial epithelium, anterior the gonad primordium. The budlet then grows and arches anteriorly relative to its parental bud (stage A2/B1), forming a transient, double-vesicle structure still connected with a stalk to the peribranchial epithelium (stage B2). As development progresses, epithelial folds reorganize the budlet into a more compartmentalized form (stage C1), eventually leading to a structure with distinct heart, gut, and neural rudiments (stage C2). In stage D, the budlet rotates to align with the primary buds and zooids, adopting a morphology similar to the adult. Organogenesis and growth continue until the budlet reaches the zooid stage.

For the budding dataset, samples selected for dissociation (either stages A1 to C2 buds with budlets or stage D budlets, Fig. 1A, B) were microdissected along with their surrounding tunic and some ampullae using fine syringe needles (30G, Microlance 3). Dissections were carried out in 5µm-filtered sea water maintained between 12-15°C, with frequent water replacement. To minimize the time between dissection and fixation, each dissociation batch included no more than 10-12 buds/budlets. Particular care was taken to avoid damaging the budding tissues during dissection to limit cellular stress. Multiple rounds of dissection were performed within the same day to increase the number of collected cells. Dissected samples were transferred to a glass 9-well plate cleaned with RNase-AWAY (Fisher) and washed twice in in calcium- and magnesium-free artificial sea water (CMFASW; 31g/L NaCl, 0.8g/L KCl, 0.29g/L NaHCO3, 1.6g/L Na2SO4, pH 8.0) with 40 U/ml RNAsin Plus (Promega). Samples were then incubated in 0.2µm-filtered 1% Trypsin (Sigma, T-9201) in CMFASW with 40 U/ml RNAsin Plus for 20-30 min. During enzymatic digestion, the tunic was separated from the tissues with needles, then tissues were mechanically dissociated by vigorously pipetting with a p200 pipet. Cell suspensions were then transferred into 2mL Protein-LoBind tubes (VWR) kept on ice. A small aliquot was taken for quality control (QC), while the remainder was immediately fixed in ACME (10% acetic acid, 15% methanol, 10% glycerol in molecular grade water (33)) with 40 U/ml RNAsin Plus for 1h on ice. The QC aliquot was stained with Erythrosin B (0.25 mg/ml in CMFASW final concentration) and counted using a Neubauer chamber grid to assess cell viability and concentration. Only preparations with over 85% cell viability were retained for further processing. Following fixation, samples were stored at −20°C for up to a few days. Fixative was removed by diluting the samples in 0.2µm-filtered 1% BSA (Sigma, A-7906) in PBS, followed by centrifugation (5min at 800g, 4°C). After carefully removing the supernatant, samples from the same dissociation batch were pooled together during resuspension in 1% BSA in PBS with 40 U/ml RNAsin Plus. DMSO was then added to a final concentration of 10% for cryopreservation, and samples were stored at −80°C for up to a month.

For the circulating cells samples, colonies at stages A1 or B2 were cleaned a couple hours before dissections. A Kim wipe was used to wipe slides, and a small paintbrush and toothpick were used under a dissecting microscope to remove biofilms, algae, parasites, dead tunic sections, and other debris near the colony. Any regions of the colony with decay were removed with a razor blade. Finally, colonies were dipped in deionized water for a few seconds to lyse any remaining bacteria and parasites. Cleaned colonies were kept in 0.22 µm filtered ASW until blood collection. Slides were dried gently with a Kim wipe. Between 100 and 200 µL of PBS-M-DTT (0.7 M D-mannitol [Sigma M9647] in 1X PBS [Gibco 70011-044] with 0.01 M dithiothreitol [DTT, Invitrogen Y00147]; DTT added fresh each day) was added with a p200 pipette to the dried colony. Multiple cuts were made with a feather scalpel in vascularized regions of the colony, including along ampullae as well as regions where vessels existed in the absence of zooids. Small bent-tipped forceps or a pipette tip were used to press gently on the top of the vessels and ampullae to extrude the blood from the cut vessels. The PBS-M-DTT with suspended blood cells was removed with a pipette and kept in a microcentrifuge tube on ice.

### 10X Chromium emulsion, library preparation and sequencing

For the budding datasets, cryopreserved samples were slowly thawed on ice, then centrifuged (5min at 800g, 4°C) to wash the DMSO. They were then resuspended in 1% BSA-PBS with 40 U/ml RNAsin Plus, filtered with a FlowMi 40µm cell strainer (Scienceware) to remove largest aggregates, and stained with 1:300 Draq5 solution (Thermo Fisher, stock 5 mM). They were then sorted by FACS to select only single cells (Supp. Fig. 1) on a BD Influx cell sorter with 4-way purity by the IPMC cell cytometry platform (Institut de Pharmacologie Moléculaire et Cellulaire – Université Côte d’Azur). Around 75k to 100k cells were collected from the FACS. The sorted cells were concentrated to around 1000 cells/µl by centrifugation. An aliquot was stained with Hoechst to check the sample quality and count the number of cells with a Countess 3 Automated Cell Counter (Invitrogen). Around 10,000 cells/sample were loaded on 10X Genomics Chromium Single Cell 3’ v3 (10X Genomics 1000123, 1000127, and 1000190), and cDNA libraries were produced following the manufacturer instructions. The libraries were then sequenced (paired-end) on an Illumina NextSeq2000 with read1=28bp (barcode + UMI) and read2 = 100bp (mRNA) excluding library index, to a depth of 200-500 million reads. When necessary, resequencing was performed to achieve a sequencing saturation of approximately 40% or higher. Library quality was checked with a BioAnalyzer. Individual 10X sample libraries were demultiplexed using Cell Ranger Makefastq v7.0.0 with default settings

For the circulating cells datasets, blood cells were pooled to generate two scRNA-seq libraries: 5 colonies for the A1 sample, and 6 colonies for the B2 sample. Each cell suspension was filtered through a 40µm strainer into a microcentrifuge tube which had been coated with 0.5mL of 10% (v./v.) bovine serum albumin (BSA, Sigma 9048-46-8) in Dulbecco’s phosphate-buffered saline (DPBS, Gibco 14200075). Cells were centrifuged at 400g for 10 min at 4°C in a swinging bucket centrifuge (Eppendorf Centrifuge 5810R). The supernatant was removed, and cells were resuspended in PBS-M-DTT. Cells were counted on a LUNA-FL Automated Fluorescence Cell Counter (Logos Biosystems L20001) using an ultra-low fluorescence slide (Logos Biosystems L12005). The cell viability prior to encapsulation was 79% (A1 sample) and 80% (B2 sample). Cell encapsulation, RT, cDNA amplification, and library preparation were carried out using the Chromium Next GEM Single Cell 3′ v3.1 Kits (10X Genomics 1000123, 1000127, and 1000190). The manufacturer’s guidelines were followed except for one step. To prepare the RT mix prior to cell encapsulation, the manufacturer protocol instructs to add nuclease-free water, then to add the cell suspension; we added PBS-M instead of nuclease-free water. Libraries were sequenced with Illumina NovaSeq 6000.

### Preparation of the gene count matrix and dimensionality reduction of the single cell datasets

The *Botryllus schlosseri* reference genome was prepared by concatenating the nuclear genome (34), with the addition of a manually annotated Piwi2 gene (Boschl.Bs14.g18242nested) and published mitochondrial genome (sc6a-b isolate, GenBank: HF548551.1, filtered to only the protein coding genes). Reads were quality-checked using FASTQC, and mapped with CellRanger count (v7.1.0 for budding datasets, v6.1.2 for circulating cells datasets). We used CellRanger automatic filtering of empty droplets for the budding samples. For the circulating cells samples, empty droplets were manually filtered by applying a soft threshold at the elbow of the barcode rank plot (retaining barcodes with nUMI > 200, Supp. Fig. 2A).

The single cell analysis was primarily conducted using Seurat (v5.1.0), complemented with selected packages from the Bioconductor (v3.18) and scanPy (v1.9.8) suites. Doublets were detected using scDblFinder (v1.16.0) and filtered out (for the “budding” datasets only). Low-quality cells were removed based on thresholds for low UMI number, low gene counts, and high mitochondrial reads percentage. Additionally, high-UMI cells were excluded from the “circulating cells” datasets to avoid potential multiplets (Supp. Fig. 2, 3).

Datasets were analyzed jointly, as separate layers within the same Seurat object but were not integrated. Normalization and identification of highly variable genes (HVGs) were performed independently for each sample, after which a consensus of HVGs was generated. Following gene expression scaling, PCA was used for dimensionality reduction. A K-Nearest Neighbors (KNN) graph was constructed, and clustering was computed at several resolutions (0.5, 1.0 and 1.5). A Uniform Manifold Approximation and Projection (UMAP) projection was computed with multiple parameters sets to ensure an optimal visualization layout. Differentially expressed genes were calculated for each cluster using Seurat built-in function FindAllMarkers. Cluster-specific markers were chosen based on high average fold changes (avg_log2FC), high proportion of expressing cells (pct.1) within the cluster and low proportion of expressing cells in other clusters (pct.2). The parameters used for each analysis will be available in the accompanying codes upon publication.

When required, subsets of clusters were reanalyzed from the normalization step to gain high resolution insights. For the analyses of endostyle cells, clusters 12 and 15 of the full budding dataset (resolution 0.5) were subset and reanalyzed using a similar pipeline. One cluster was identified as low quality (characterized by low UMI and gene counts, and absence of clear marker genes) and was removed before a final round of reanalysis.

For the comparison between the circulating cells dataset and the budding dataset, quality checks and preprocessing were performed on each sample separately, as described above. All samples were analyzed jointly, without integration, using a similar pipeline as described above. A clustering for this joint dataset was calculated, at resolution 0.5. Circulating status of each cluster was assigned based on the proportions of cells from the circulating cells dataset. Clusters with <1% of cells, between 1 and 10%, and >10% from the circulating cells dataset were annotated as “non circulating”, “rarely circulating” or “frequently circulating”, respectively. These joint clusters and the circulation status annotation were transferred to the budding dataset-only UMAP based on the cell barcodes.

For the specific analyses of hematopoietic-lineage hemocytes, cells from the PAGA module “hemocytes” were subsetted and analyzed with Seurat. A new UMAP projection was generated using scanPy by recomputing HVGs, PCA, and constructing a KNN graph. The resulting UMAP coordinates were then transferred back to the Seurat object for downstream analyses.

For the “budding tissues” subclustering, we excluded cells assigned to the “hemocytes” and “phagocytes” PAGA modules, and analyzed the remaining subset. This revealed a low-quality cluster (cluster 24 in “noBlood_clusters_0.8” of the budding tissues subclustering), which was subsequently removed. We were then able to resolve and separate branchial from peribranchial epithelial cells and identify the budlet population more clearly.

For the “budlet-specific” subclustering, we further refined the analysis by selecting cells from the “budding tissues” subclustering (resolution 0.8), specifically those belonging to early peribranchial epithelium (cluster 2), early branchial epithelium (cluster 9), budlet stalk septum cells (cluster 30), and *Bhf+* cells (cluster 31). This subset was reanalyzed using the same basic pipeline as for the full dataset. Marker genes for the subclustering were identified at a clustering resolution of 0.4.

Functional annotation was carried out using Eggnog emapper (v2.1.10) with default parameters to assign GO terms to genes. Differentially expressed genes (DEGs) for selected clusters or cluster groups were identified using the FindMarkers function in Seurat, comparing each cluster against the rest of the dataset. GO term enrichment analysis was performed with topGO (v2.54.0): for the body wall muscles, heart and germline clusters, up- and down-regulated genes were used to compute enrichment with the Kolmogorov-Smirnov test; for the putative phagocyte, hyaline amoebocyte, and hemoblast clusters enrichment was calculated on up-regulated genes only using Fisher’s test.

### Potency inference, trajectory inference

To infer cell potency, we used CytoTRACE (v0.3.3, (35)) on both the entire dataset and on selected subsets. For trajectory inference, we applied different complementary approaches. PAGA (Partition-based Graph Abstraction, (36)) was used to assess the connectivity between clusters, revealing relationships between transcriptionally similar populations, which can reflect differentiation trajectories. For analysis on the full budding dataset, we removed lowest quality cells (number of UMI<400), as they tend to skew the connections. For each PAGA computation, we tested different thresholds to remove weak cluster connections.

To compute RNA velocity, we demultiplexed spliced and unspliced count matrices by running Velocyto (v0.17, (37)) on the CellRanger outputs. RNA velocity vectors were then calculated with the scVelo dynamic model (v0.3.2, (38)) using a scanPy object converted from the Seurat reference. In parallel, Monocle 3 (v1.3.7, (39)) was used to infer putative differentiation trajectories based on transcriptome similarity.

### Immunofluorescence and *IN SITU* hybridization by WMISH and HCR

Proliferating cells were revealed by fluorescent immunostaining for phospho-histone 3, following the protocol in (40), and additional phalloidin staining was performed according to (12).

Gene expression patterns were examined using two methods. Whole mount fluorescent *in situ* hybridization (WMISH), was performed using the protocol described in Langenbacher et al. 2005 (32) REF. For Hybridization Chain Reaction (HCR), probes were designed using Özpolat Lab’s HCR *in situ* probe generator (41). Samples were fixed, stored, and dissected, and post-fixed following the WMISH protocol. The remaining steps followed the protocol described in (42) from the pre-hybridization stage, with some modifications. Following post-fixation washes, samples were washed twice in 2xSSC and once in 50% 2xSSC - 50% probe hybridization buffer(32). Samples were pre-hybridized for 4 h at 37°C in hybridization buffer. The pre-hybridization solution was removed and replaced by 1µM of probes in hybridization solution, let to hybridize for one night to 3 days at 37°C. Samples were amplified with 60nmol/L each hairpin (h1 and h2) conjugated with an Alexa 488 or 647 (B1 or B2 hairpins, Molecular Instruments) and incubated one night to 2 days at 23°C. Hoechst 33342 staining was performed either during the amplification step, or after amplification washes at 10µg/mL, for 30min. As negative controls, we either used no probes, or probes designed for other species, which yielded similar results with low aspecific background and some autofluorescent cell types. Primers and probes sequences for WMISH are listed in Annex 1. HCR probes sequences are available upon request.

Samples were mounted in 75% glycerol or Vectashield Plus (Vector) and imaged using a Leica SP8 or Leica Stellaris confocal microscope. Images were processed and analyzed with ImageJ.

## Results

### 1. A MULTI-STAGE ATLAS OF *B. SCHLOSSERI* PERIBRANCHIAL BUDDING

To investigate the molecular and cellular changes during *B. schlosseri* asexual development via peribranchial budding (1), we collected buds and their associated budlets from stage A1 to stage C2 and budlets at stage D (Fig. 1A-B, Supp. Fig. 1A), all from an isogenic colony (*i.e.* derived from a single, founder larva). Following dissociation, single cells were sorted by FACS and loaded onto 10X Chromium V3 chips (Supp. Fig. 1B). Cell viability and RNA quality were confirmed (Supp. Fig. 1A, C). Sequenced reads were mapped on the latest version of *B. schlosseri* genome (34). Mapping metrics are reported in Supp. Tab. 2, and confirm overall good quality datasets. CellRanger initially retained 78,257 cells over all the samples, which were subsequently filtered to remove lower-quality cells and doublets using soft thresholds (Supp. Fig. 2 and 3), resulting in a final budding dataset of 66,227 cells. All samples were preprocessed and analyzed jointly.

Clustering resulted in 42 transcriptionally distinct clusters, visualized using Uniform Manifold Approximation and Projection (UMAP, Fig. 1C). Quality metrics across the full dataset and within clusters are shown in Supp. Fig. 4.

Each cluster comprises cells from multiple budding stages (Fig. 1D, Supp. Fig. 5A) and most of the clusters display a clear gradient of stages along the UMAP (Fig. 1D), which is consistent with the progressive state of development of the bud and budlet and strongly point out potential differentiation trajectories within the dataset. Importantly, we chose not to integrate the samples, as stage-specific biological variation is central to our analyses. In addition, the two biological replicates for stage B2 overlapped well, indicating that sample integration was not necessary (Supp. Fig. 5B).

To explore possible differentiation trajectories, we applied CytoTRACE and RNA velocity analyses. Although CytoTRACE is generally considered less reliable for multiple heterogeneous tissues (35, 43), its analysis of the entire dataset supports the presence of cellular differentiation dynamics, consistent with the developmental progression captured (Supp. Fig. 6A-C). Although only 5% of the genes were considered unspliced, limiting the resolution, RNA velocity vectors projected on the UMAP are also globally consistent with the differentiation gradient suggested by CytoTRACE and the developmental staging gradient, with some exceptions (Supp. Fig. 6D).

To further explore relationships between clusters, we computed PAGA (Partition-based Graph Abstraction, (36)) on the underlying KNN graph, revealing nine major groups, (Fig. 1E, Supp. Fig. 6E). Differential expression analyses show that clusters have distinct transcriptomic signatures (Fig. 1F). This allowed us to identify cluster-specific marker genes and annotate the cluster’s identity. Initial validation was guided by previously published gene expression data from *B. schlosseri* (Supp. Tab. 3) and related tunicate cell-type literature. In several cases, known gene expression patterns were sufficiently specific to allow direct annotation of certain clusters.

Cluster 7 and 10 express *Myh3* (*Boschl.Bs15.g19623-Myh6*) a marker of body wall muscles, and *Ebf* (*Boschl.Bs10.g14018-EBF1*), which is specific of developing muscles and nervous system (Supp. Tab. 3, Supp. Fig. 7A, (23)). These clusters also express myogenic markers such as *Mrf (Boschl.Bs4.g5757-MYOD1)*, *Troponin-like* (Boschl.Bs10.g14218-TNNT2.1) and a *Titin-like (Boschl.Bs5.g6251)* (44), consistent with muscle lineage identity. Several terms associated with muscle identity are highly enriched in this cluster, including “skeletal myofibril assembly” or “sarcomere organization” (Supp. Fig. 7B).

Cluster 24 was annotated as heart cells based on the expression of *Myh2 (Boschl.Bs8.g10861-MYH3.1)*. This cluster also expresses transcription factors known to be involved in heart development, including *Gata-a, Tbx1/10, FoxF* and *Nk4* (Supp. Tab. 3, Supp. Fig. 7A, (11, 23)). Similarly, GO terms associated with muscular identity, and in particular with cardiac muscle, are enriched in this cluster (Supp. Fig. 7B).

Cluster 3 was identified as germline cells based on the strong expression of *Vasa (Boschl.Bs4.g5801-DDX4),* a well-established germline marker, as well as *Piwi1 (Boschl.Bs5.g6728-PIWIL1.1)* (Supp. Tab. 3, Supp. Fig. 7A, (32)), along with the expression of multiple genes from the germline multipotency program (GMP) genes (see below, (45)). Accordingly, several GO terms associated with meiosis are highly enriched in this cluster (Supp. Fig. 7B).

Clusters 6, 13 and 14 showed enriched expression of *Tgf-ꞵ-f (Boschl.Bs8.g10511),* which has been described in follicular and germline support cells (Supp. Tab. 3, Supp. Fig. 7A, (32)). Given the heterogeneity and transcriptomic complexity of these clusters, they were conservatively annotated as *Tgf-ꞵ-f+* cells.

To elucidate unknown cluster identities, we performed whole-mount fluorescent *in situ* hybridization and successfully implemented a hybridization chain reaction (HCR) protocol tailored for *B. schlosseri* (see Methods). For robust cell-type annotation, we defined marker genes as those exhibiting both strong differential expression and high specificity within a given cluster. To minimize false assignment, we selected at least two marker genes per cluster.

Clusters 0, 2 and 20 were annotated as the branchial and peribranchial epithelia (excluding the endostyle and stigmata), based on the expression of the collagen *Boschl.Bs16.g20741-COL2A1* (Fig. 2A), as well as *Pax1/9 (Boschl.Bs6.g8552-PAX9.1,* Fig. 2B) or *Pax2/5/8a (Boschl.Bs16.g20739-PAX5.1,* Fig.2C) for the branchial and peribranchial epithelia respectively. These clusters also express transcription factors known to be expressed in these tissues, such as *Gata-b*, *Tbx1/10*, *Tbx2/3* and *Nk4* (Supp. Tab. 3, Supp. Fig. 7A, (11, 23)).

**Figure 2.**
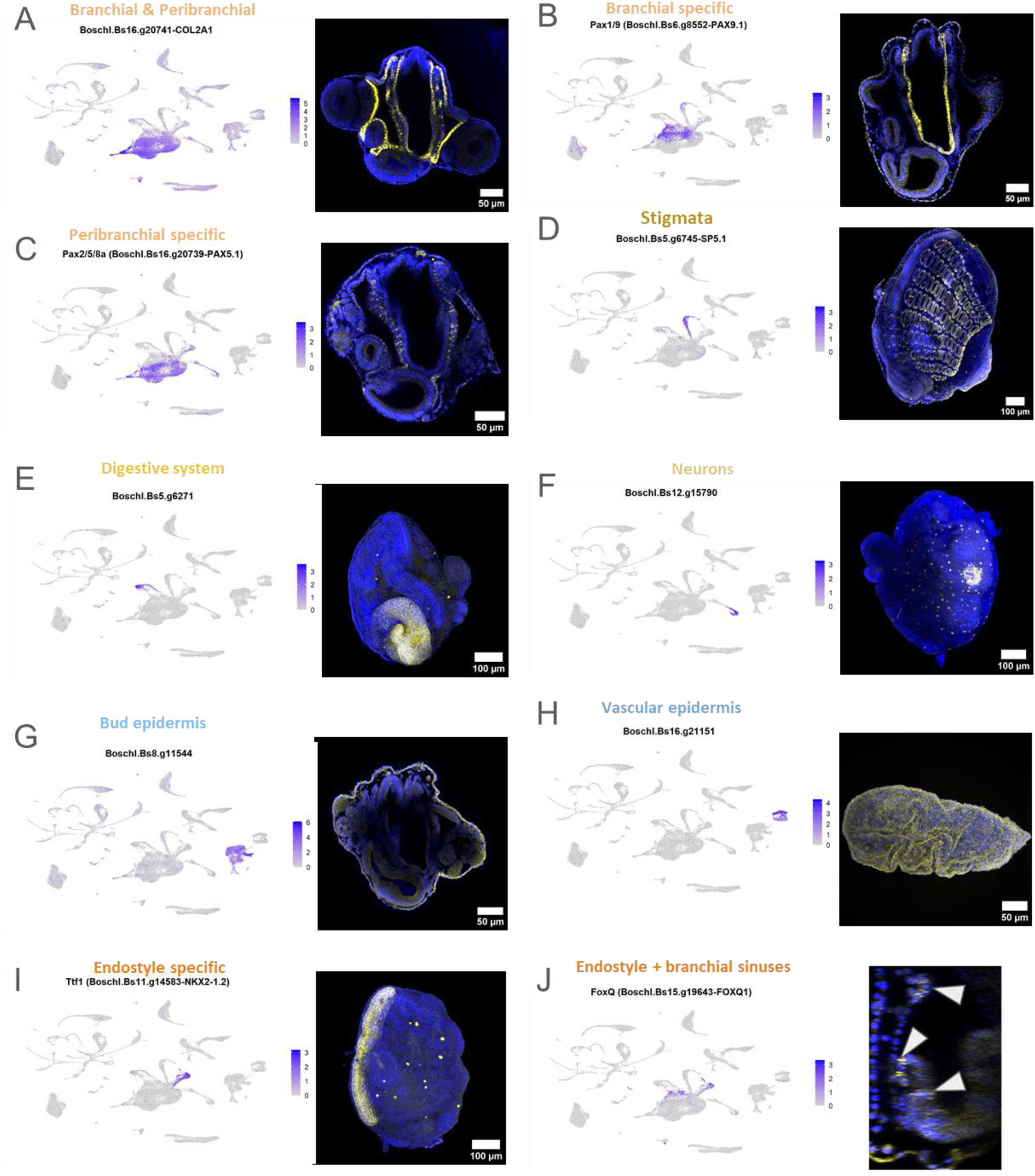
Annotation of the budding tissues clusters from gene expression patterns. **(A-J)** Expression plots (on the left) and WMISH or HCR confocal images (on the right) of marker genes. The probe signal is in yellow and the Hoechst staining is in blue. **(A)** *Boschl.Bs16.g20741-COL2A1*, stage A1 bud. **(B)** *Pax1/9 (Boschl.Bs6.g8552-PAX9.1)*, stage B1 bud. **(C)** *Pax2/5/8a (Boschl.Bs16.g20739-PAX5.1)*, stage A1 bud. **(D)** *Boschl.Bs5.g6745-SP5.1*, stage C1 bud. **(E)** *Boschl.Bs5.g6271*, stage B1 bud. **(F)** *Boschl.Bs12.g15790,* stage B1 bud. **(G)** *Boschl.Bs8.g11544*, stage A1 bud. **(H)** *Boschl.Bs16.g21151*, ampulla. **(I)** *Ttf1 (Boschl.Bs11.g14583-NKX2-1.2)*, stage C1 bud. **(J)** *FoxQ* (*Boschl.Bs15.g19643-FOXQ1*), stage C1 bud endostyle, expression zones are pointed with arrowhead.

Cluster 9 was identified as the stigmata (peribranchial side), characterized by the expression of *Boschl.Bs5.g6745-SP5.1* (Fig. 2D) and *Boschl.Bs15.g20397* (Supp. Fig. 8), as well as FoxA1 (Supp. Tab. 3, Supp. Fig. 7A, (11)).

Cluster 18 corresponds to the digestive system, as shown by the expression of the marker genes *Boschl.Bs5.g6271* (Fig. 2E) and the annexin *Boschl.Bs8.g11037-ANXA13.2* (Supp. Fig. 8). This cluster also expresses typical markers of the digestive system including *Gata-a, FoxA1* and *FoxA2* (Supp. Tab. 3, Supp. Fig. 7A, (11)).

Cluster 17 was annotated as the nervous system, based on the expression of *Boschl.Bs12.g15790* (Fig.2 F) and a tubulin-α (*Boschl.Bs8.g11041-TUBA3D.2)* (Supp. Fig. 8), although the latter is also detected in developing oocytes, which are not present in this dataset. This cluster also expresses markers of the nervous system, such as *Zic.r-a* (Supp. Tab. 3, Supp. Fig. 7A, (22)).

Clusters 4, 11 and 32 correspond to epidermal cells, while cluster 31 represents the internal siphon epithelium. These are all marked by *SoxB2* (*Boschl.Bs14.g18200-SOX14.1*, Supp. Fig. 8) and one of the two cellulose synthase genes (*Boschl.Bs9.g12156*, homologous of *Ciona CesA*, Supp. Fig. 7A). Cluster 4 and 32 represents epidermis of the bud and budlet, based on the expression of *Boschl.Bs8.g11544* (Fig. 2G) and *Boschl.Bs10.g13389* (Supp. Fig. 8). Cluster 11 corresponds to the vascular epidermis, expressing *Boschl.Bs16.g21151* (Fig. 2H) and *Boschl.Bs9.g12759-SEMA5A.3* (Supp. Fig. 8). Cluster 31 expresses *Boschl.Bs1.g1247* (Supp. Fig. 8), marking the internal epithelium of the siphons.

Cluster 12 and 15 correspond to the endostyle and adjacent regions in the branchial epithelium. *Boschl.Bs11.g14583-NKX2-1.2* and *Boschl.Bs5.g6849* are specifically enriched in cluster 14, and both are restricted to the endostyle (Fig. 2I, Supp. Fig. 8). *Boschl.Bs11.g14583-NKX2-1.2* is potentially orthologous to *Ttf1* from *Ciona intestinalis*, a known endostyle marker (46–48). *FoxQ* (*Boschl.Bs15.g19643-FOXQ1*) is specific to clusters 12 and 15, and labels the part of the endostyle, its flanking regions and part of the branchial epithelium (Fig. 2J).

### 2. Assessing endostyle regionalization and putative stem cell niche role

The endostyle in tunicates is a multifunctional pharyngeal structure homologous to the vertebrate thyroid gland (48), and potentially to other vertebrate pharyngeal derivatives (49). In *Botryllus schlosseri*, the endostyle and cells in the sub-endostylar region have also been proposed to serve as a niche for putative stem cells involved in bud development (26).

Therefore, to investigate the molecular and cellular heterogeneity of the endostyle cells in our dataset, we subsetted and reanalyzed cells from clusters 12 and 15 (Fig. 3A). The tunicate endostyle exhibits a highly organized structure, with distinct zones that vary in cellular morphology. These zones reflect specialized functions, such as mucus secretion, iodine metabolism, and ciliary movement, each associated with specific cellular characteristics. (Fig. 3B, (49, 49, 50)). This subclustering revealed 12 transcriptionally distinct cell clusters. One cluster (cluster 0) was predominantly composed of cells from early stages of endostyle formation, i.e. budlet stage D and bud stage A1, while the remaining clusters corresponded to latter stages of development (A2 to C2). As previously observed in the global UMAP, the arrangement of cells within clusters reflects the developmental progression, strongly suggesting these clusters represent differentiation trajectories (Fig. 3A).

**Figure 3.**
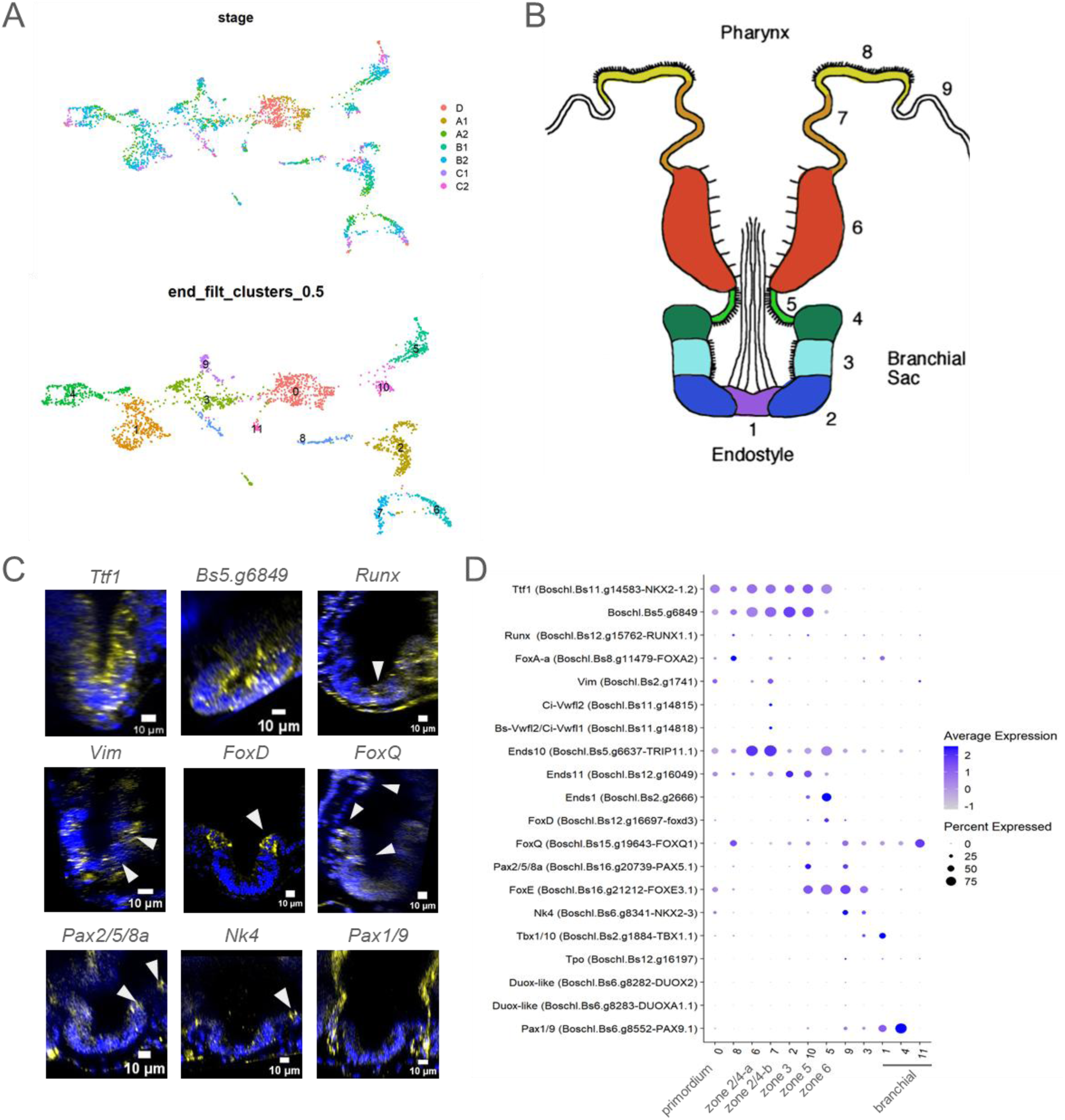
Sub-clustering of the endostyle cells and spatial regionalization. **(A)** UMAP projection of the subclustered endostyle cells, colored by stage (top panel) and by cluster (lower panel) at resolution 0.5. **(B)** Schematic representation of the endostyle in transverse section showing the eight cell bands (modified from Satoh et al., 2007). **(C)** WMISH or HCR of genes expressed in the endostyle (orthogonal projections of confocal z-stack images); white arrowheads point at transcript expression areas **(D)** Scaled expression of known markers for ascidians endostyle zones, and the inferred correspondence with UMAP clusters (panel A).

To assign identities to each cluster, we combined known expression patterns from *B. schlosseri* and *Ciona sp*. with novel data from WMISH and HCR (Fig. 3C, D, Supp. Tab. 4). This integrative approach allowed us to propose a correspondence between transcriptomic clusters and histologically defined endostyle zones (Fig. 3D).

Subclusters 6 and 7 likely represent populations of zones 2 and 4, although it remains unclear whether the two clusters correspond to distinct zones or represent shared cell types present in both regions. This reflects the unclear orthology between Von Willebrand Factor genes in *Ciona* and *Botryllus*, which serve as regional markers in the former (Fig. 3D, Supp. Tab. 4) and underscores the need for more precise spatial validation.

Although *Tpo* and *Duox,* genes classically associated with zone 7 (51, 52), were minimally expressed in our datasets (likely due to the early developmental stages sampled, Fig. 3D), two *Duox* paralogs (*Boschl.Bs6.g8282* and *Boschl.Bs6.g8283*) were specifically detected in a small number of cells in subcluster 9. This suggests that subcluster 9 may correspond to zone 7. However, the precise boundaries between zones 7, 8 and 9, as well as their transitions to the surrounding branchial basket epithelium, remain unresolved. Further spatial validation using double HCR might be required to refine our understanding of these regions.

This molecular atlas of the endostyle could serve as a resource to explore the putative niche roles suggested for the endostyle in *B. schlosseri*. For instance, *Wnt* have been shown to be involved in short range signaling and crucial in regulating the niche microenvironment for a large variety of stem cells in vertebrates and invertebrates (53, 54). In our dataset, several *Wnt* are being expressed in the endostyle clusters, in particular in the ones corresponding to zone 5 (subcluster 10) and 6 (subcluster 5, Supp. Fig. 9).

### 3. scRNA-seq analyses reveals no distinct population of pluri-/multipotent stem cell during budding

Adult pluripotent or multipotent stem cells underpin various forms of non-embryonic development, namely agametic reproduction and whole-body regeneration, in a wide range of animals such as acoels, annelids, cnidarians, and sponges (45, 55–57). In these organisms, adult stem cells proliferate and are molecularly characterized by the expression of multiple genes associated with the Germline Multipotency Program (GMP, (45)). In tunicates, genes associated with the GMP, such as *Piwi-2*, *Vasa* and *Pl10*, have been proposed as potential markers of pluripotent or multipotent cells involved in agametic development and whole-body regeneration (18, 27, 29). Indeed, *Piwi-2* knock-down disrupts vascular budding in *Botrylloides leachii* (18, 27), and both *Piwi-2* and *Vasa* are highly expressed in proliferating cells during the early stages of vascular budding in *B. diegensis* (29). In contrast, during peribranchial budding in *Botryllus schlosseri,* the role of *Piwi* and *Vasa* appears to the confined to the germline (27, 58). Similarly, in *Botryllus primigenus* peribranchial budding *Piwi-2* and *Vasa* expression are restricted to the germline (59).

To identify potential multi- or pluripotent candidate stem cells involved in budding, we first examined which clusters might contain actively proliferating cells. Most clusters showed expression of proliferative markers (Supp. Fig. 10A), consistent with the active growth and morphogenesis of bud and budlet tissues. This was further supported by immunofluorescence staining for phospho-histone H3 (Supp. Fig. 10B). While this broad proliferative signal aligns with the expected cellular dynamics of budding, it does not permit the identification of discrete proliferating cell populations, even when comparing different budding stages.

When we screened for cells co-expressing GMP markers, we found enrichment in cluster 4, which corresponds to the germline lineage (Supp. Fig. 11A). Other clusters, including the budding tissues in formation, express a subset of the GMP genes. Although we cannot completely exclude the presence of a small population of GMP-positive cells outside this cluster, no such cells were detected in our dataset.

In *B. schlosseri*, the co-expression of several paralogs of Yamanaka factors (in human: *Sox2*, *c-Myc*, and *Pou5f1)* —particularly *SoxB1*, *Myc*, and *Pou3*— has been proposed to be associated with candidate stem cells in adults (60). Another paralog of the Yamanaka factor *Pou5f1 - Pou2-* has been described as restricted to early budlet and the female germline (60). In the reference genome used to map our scRNA-seq reads (34), the *Pou3* gene is missing, while *Pou2* (Boschl.Bs11.g15669-POU2F1.1) appears to be ubiquitously expressed, with relatively higher expression in circulating cells, specifically clusters 5, 27, and 40, which may correspond to amebocytes cell populations ((61); see below). Cells co-expressing *SoxB1* (Boschl.Bs11.g14812-SOX3.1) and *Myc* (Boschl.Bs10.g13375) were found in clusters 9, 12 and 15, which correspond to the stigmata, a portion of the branchial epithelium, and the endostyle, respectively (Supp. Fig. 11B). This is consistent with the described expression in tissues other than candidate stem cells in bud and budlets (60). However, their distribution did not suggest the presence of a distinct pluripotent population nor specific links to the origin of the budlet clusters (see below).

### 4. Circulating mesenchymal cells reveal the diversity and differentiation trajectories of a complex hematopoietic system

*B. schlosseri* displays a large diversity of circulating cells within its internal and extracorporeal vascular system (62–64). Given that our scRNAseq datasets were derived from whole buds and budlets, including some of the surrounding vasculature, we hypothesized that circulating cells were captured in the budding datasets. This was supported by the presence of vascular epithelial cells (cluster 11, Fig. 1C, Fig. 2H).

To assess which clusters might correspond to circulating cell types, we generated two scRNAseq datasets from cells directly isolated from the vasculature by bleeding out adult colonies of *B. schlosseri.* These cells were immediately processed for scRNAseq without any dissociation nor sorting or enriching step. Quality control for these circulating cells datasets is detailed in Supp. Fig. 2 and Methods.

Joint analyses of the budding datasets and circulating cells datasets revealed that only a subset of clusters contained cells from the circulating cells datasets (Fig. 4A Supp. Fig. 12A, B). Clusters with less than 1% of circulating dataset-derived cells were classified as non-circulating, likely representing epithelial or non-mobile mesenchymal populations (Supp. Fig. 12C). The remaining clusters, which contained at least 1% of circulating cells, were considered to represent cell types contributing to the circulatory pool. Among them, clusters with 1-10% of circulating cells likely reflect mesenchymal cells that are mostly resident in bud, budlets or ampullae but rarely, or transiently circulate (Supp. Fig. 12C), while clusters containing more than 10% of circulating cells were considered as frequently circulating cells (Supp. Fig. 12C). Joint clustering, circulating status, and their mapping on the reference UMAP projection (full budding dataset) are shown in Supp. Fig. 12B, D.

**Figure 4.**
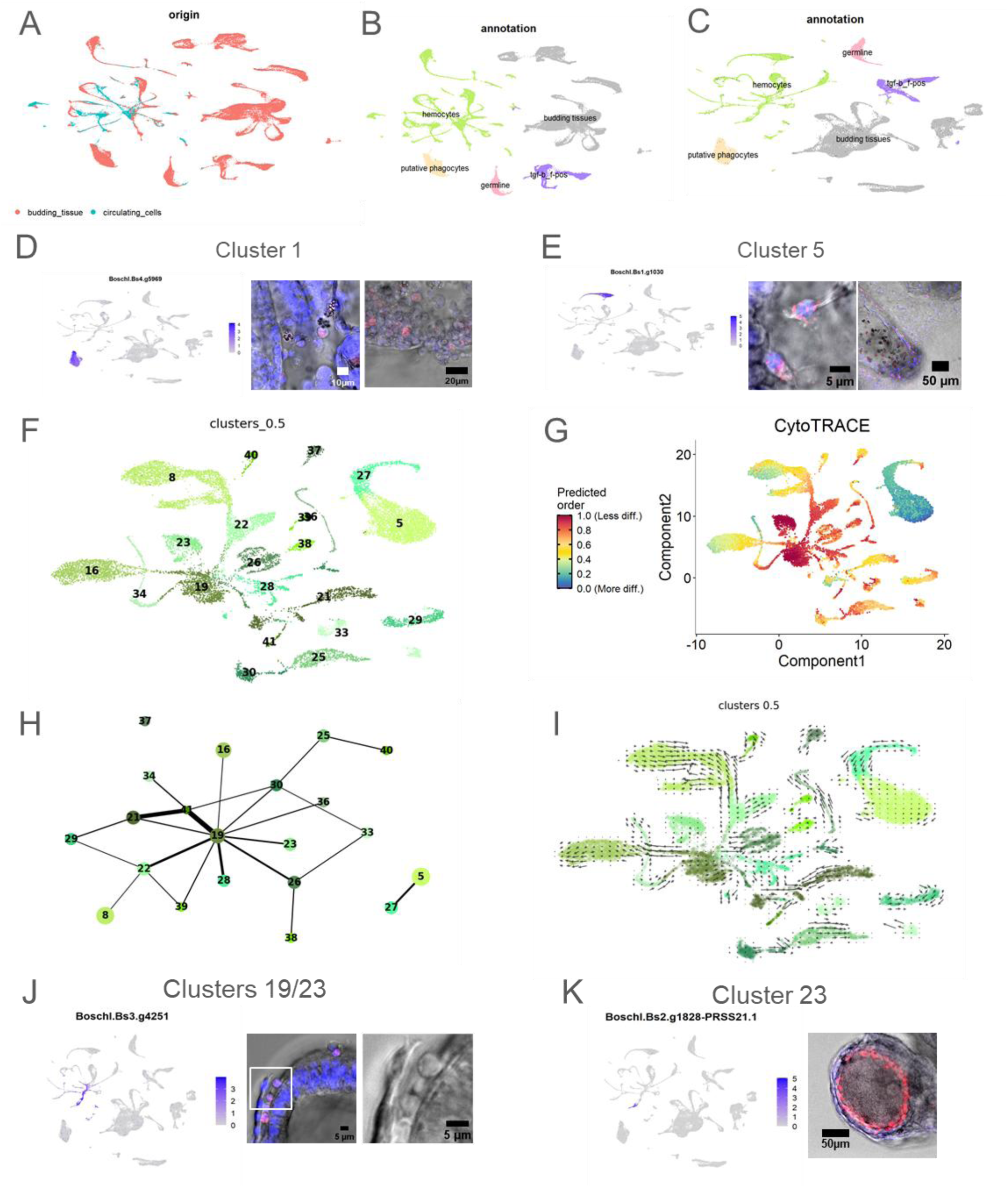
Diversity of circulating cells and differentiation trajectory inference. **(A)** UMAP projection of the joint analysis of the full budding (red) and the circulating cells (blue) datasets. **(B-C)** PAGA modules of circulating cells, represented with **(B)** same UMAP as in (A) with joint datasets and with **(C)** UMAP projection of the full budding dataset; colors in (B) and (C) according to the 4 corresponding PAGA groups in Fig 1E. **(D-E)** Expression plots and HCR of a marker gene of the **(D)** putative phagocytes cluster and **(E)** putative hyaline amoebocytes clusters. **(F-I)** Separate analysis of the hemocyte module. **(F)** UMAP projection with cluster numbers corresponding to the full budding dataset (resolution 0.5, see Fig 1E) and **(G)** corresponding CytoTRACE score, **(H)** abstracted graph (PAGA), **(I)** projected RNA velocity vectors. **(J-K)** Expression plots and HCR of marker genes of **(J)** putative hemoblasts (cluster 19) and resident hemoblast-like (cluster 23), or **(K)** resident hemoblast-like only (cluster 23). For HCR pictures: the probe signal is in red, Hoechst staining in blue, and the transPMT in grey.

PAGA analyses of the full budding dataset showed that the circulating cell clusters grouped into four distinct, minimally connected PAGA modules (Fig. 1E, Fig. 4B, C), suggesting that either intermediate cell states were not captured in our datasets or, more plausibly, that these clusters belong to separate lineages.

These observations align with previous studies proposing that *B. schlosseri* harbors a complex vertebrate-like hematopoietic system. Moreover, UMAP connectivity patterns further support the notion that at least some circulating cell types are lineage-related (Fig. 1C, D; Fig. 4A).

The first two PAGA groups correspond to (1) the germline and (2) the Tgf-*ꞵ-f+* clusters and the closely associated small cluster 35 strongly expressing *Bhf* (Fig. 1E, (65)). Among Tgf-*ꞵ-f+* cells, support cells are associated with immature germ cells that circulate during stage B2 to colonize newly forming budlets (17, 66). Only one of the Tgf-*ꞵ-f+* clusters was represented in the circulating cells, and as “rarely circulating” (Fig. 4B, C; Supp. Fig. 12B, D).

A third group includes a single large “rarely circulating” cluster. HCR on its marker gene *Boschl.Bs4.g5969* revealed expression in cells of variable morphology, some of which are at least 10µm of diameter, and are reminiscent of described phagocyte morphology (Fig. 4D, (67, 68)). Negative controls are shown in Supp. Fig. 14A. These cells are likely to be mostly resident in tissues, as they are present in much higher numbers in our budding datasets than in the circulating datasets (Supp. Fig. 12B, D). GO terms enriched in this cluster include terms related to innate immune response, such as neutrophil degranulation (GO:0043312) and terms related to migration and localization of the cell (GO:0051674, GO:0030335) (Supp. Fig. 13B).

The fourth and largest group includes all the “frequently circulating”, a few “rarely circulating” and one “non circulating” cluster and was annotated as hemocytes. They are highly connected in the PAGA graph, indicating that they may belong to a common hematopoietic lineage (Fig. 4B, C, Fig. 1E). To link transcriptomic data to known hemocyte morphotypes, we examined gene expression for some of these clusters. Cluster 16 expresses a phenoloxidase (*Boschl.Bs1.g975-PPO1*), a known marker for granular amoebocytes and morula cells (Supp. Fig. 14C). Cluster 29 expresses one *Botryllin* paralog (*Boschl.Bs11.g14579*, (69)), also associated with morula cells morphotype (Supp. Fig. 14C).

Cluster 5 expresses the kringle-domain containing protein *Boschl.Bs1.g1030*; HCR revealed an amoebocyte morphology with few vacuoles, consistent with hyaline amoebocyte morphotype (Fig. 4E, negative controls in Supp. Fig. 13A, (62, 67)). These cells localize both in circulation and within the tunic, especially around the ampullae (Fig. 4E). They also express a cellulose synthase (*Boschl.Bs9.g12156*), suggesting a role in tunic turnover (Supp. Fig. 13C). GO terms enriched in this cluster are diverse, and include apoptotic cell clearance (GO:0043277, Supp. Fig. 14B). Notably, this cluster, as well as in cluster 21 and 27, express *Athena (Boschl.Bs11.g14504)*, a gene that was shown to be necessary at stage D/A1 for future budding organogenesis (Supp. Fig. 13C, (70)).

To identify potential progenitors, we focused on the “hemocytes” clusters (Fig. 4B, C) and reanalyzed them separately (Fig. 4F). CytoTRACE analysis, which estimates the relative differentiation status, ranked subclusters 19 and 23 as having the highest developmental potential (Fig. 4G, Supp. Fig. 14A). These subclusters also show high expression of proliferation markers (Supp. Fig. 11A), and subcluster 19 is topologically central in the PAGA graph (Fig. 1H), consistent with a progenitor role. We further performed RNA velocity analysis using scVelo. The inferred velocity vectors supported the differentiation trajectories radiating from subclusters 19 and 23 (Fig. 1I). GO terms enriched in these clusters include terms related to the cell cycle, such as “G1/S transition of mitotic cell cycle” (GO:0000082) and hematopoiesis, such as “hematopoietic or lymphoid organ development” (GO:0048534, Supp. Fig. 15B).

Together, these analyses strongly suggest that subclusters 19 and 23 represent hematopoietic progenitor populations. Hemoblasts, previously described in *B. schlosseri* and other tunicates as putative progenitors for every type of hemocytes, are small (∼5µm), round cells, with high nuclear-to-cytoplasmic ratio and classical undifferentiated morphology (62, 67, 68). HCR on *Boschl.Bs3.g4251*, enriched in clusters 19/23 (Fig. 4J), and *Boschl.Bs3.g4360*, enriched in clusters 19/28 (Supp. Fig. 14C), confirmed their expression in cells with hemoblast-like morphology, further supporting this identity. The genes *Boschl.Bs2.g1828-PRSS21.1* and *Boschl.Bs13.g17339* enriched in cluster 23, are notably expressed in “test-like” cells surrounding the more mature oocytes (Fig. 4K, Supp. Fig. 14D), under the first layer of follicular cells. This localization is consistent with their “non circulating” status detected above (Supp. Fig. 12C, D). Negative controls are shown in Supp. Fig. 14E.

Our integrative analyses identified a broad identity of circulating cells in *B. schlosseri,* most of which are part of a common hematopoietic-like lineage, and genes that are specifically expressed in probable stem cells or progenitors of the hematopoietic lineage.

### 5. Identification of peribranchial epithelial founder cells initiating budding

To identify the earliest budlet-forming cells in our dataset, we screened for the co-expression of three genes previously reported to mark early budlet identity: the intermediate filament *Vim* (*Boschl.Bs2.g174*, (12)), and the two transcription factors *Runx (Boschl.Bs12.g15762-RUNX1.1,* (12, 32)) and *Nk4* (*Boschl.Bs6.g8341-NKX2-3*, (5, 23)). Cells expressing all three markers were restricted to a small subset, of the early branchial and peribranchial cluster 2 (Fig. 5A) ranging from stage D to stage C2, suggesting that our dataset includes secondary bud (budlet) cells.

**Figure 5.**
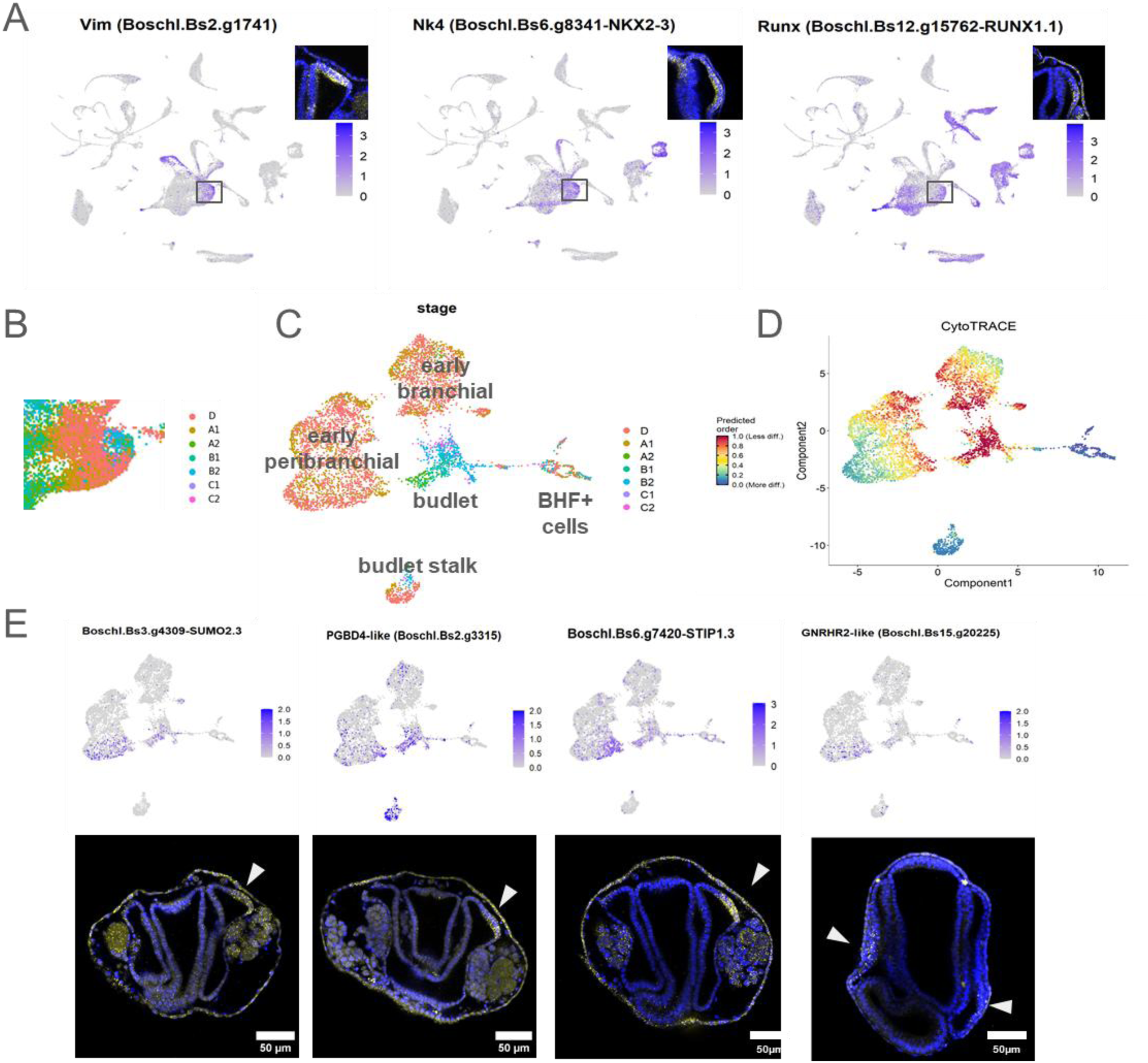
Budding founder cells in the peribranchial epithelium have a specific transcriptomic signature. **(A)** Expression plots and corresponding HCR of known budlet markers Vim, Nk4 and RUNX1 with relative HCR expression in the budlet cells population (squared). **(B)** Cropped view of the full budding dataset UMAP (Fig. 1D) showing the gradient of stages in the budlet subpopulation (squared in (A)). **(C)** UMAP projection of the re-clustered “budlet” subset, containing the budlet cluster and neighboring population clusters (subseted clusters visible in Supp. Fig. 16, 17). **(D)** CytoTRACE score of the cells in the “budlet” subset. **(E)** Expression plots and corresponding HCR of marker genes of the budding founder cells in stage C2-D budlet or A1 bud+budlet.

To increase resolution in the budding tissues, we removed most of the circulating cell clusters (retaining only germline and Tgf-ꞵ-f+/Bhf+ cells) and reanalyzed the data (Supp. Fig. 15A, C, D). In this subset, the A1-C2 budlet cells (as detected by the expression of *Vim*, *Runx* and *Nk4* and the gradient of stages, similarly to Fig.5A) emerged as a distinct subcluster (subcluster 26) that branches from the early peribranchial population (subcluster 2). PAGA analysis confirmed that this cluster is highly connected to the early peribranchial subcluster 2, as well as to the early branchial subcluster 9, to the stalk septum subcluster 30, and to subcluster 31 corresponding to the BHF+ follicular-adjacent cells (Supp. Fig. 15D). Notably, the Bhf+ cell cluster appears to form a transcriptional bridge between the budlet and the Tgf-ꞵ-f+ clusters. Additionally, both the heart (cluster 23) and the early neuronal progenitor (cluster 28) clusters show transcriptional affinity with the Tgf-ꞵ-f+ group.

Expression patterns of *Mbnl2* (*Boschl.Bs14.g18091-MBNL1*), *Vim* (*Boschl.Bs2.g174)* and *Runx (Boschl.Bs12.g15762-RUNX1.1)* in the stalk septum (Supp. Fig. 15E) revealed that this structure remains connected to the peribranchial epithelium rather than being a fold of the epidermis (Supp. Fig. 15F) as suggested by Burighel and Brunetti (71).

To better identify the cellular relationships around budlet formation, we further subsetted and reclustered these five strongly linked clusters (Fig.5C, Supp. Fig. 17A). UMAP projections revealed that the budlet cells from stage A2 (labeled as “budlet” in Fig. 5C) diverge from a distinct small population of early peribranchial cells (cluster 3 in Supp. Fig. 16B). PAGA revealed connectivity between the budlet cluster and clusters 1 and 3 from the early peribranchial cells population (Supp. Fig. 17D). Trajectory inference using Monocle3 and the staging gradient identified cluster 3 as the most probable population of origin for budlet cells (Supp. Fig. 16E). In contrast, RNA velocity analysis with scVelo produced directionalities opposite to the expected developmental progression (Supp. Fig. 16F) and was considered unreliable in that case.

To confirm spatial localization, we selected several genes enriched in putative founder cells cluster 3 and validated their expression by HCR. Genes such as *Sumo2-like (Boschl.Bs3.g4309-SUMO2.3), Pgbd4-like (Boschl.Bs2.g3315), Stip1-like (Boschl.Bs6.g7420-STIP1.3),* and *Gnrhr2-like (Boschl.Bs15.g20225)* showed specific expression in a group of cells from the peribranchial epithelium located anterior to the gonadal primordium as early as stage D (Fig.5D, Supp. Fig. 17), supporting their identity as the founder population for budlet initiation. Gene expression in this population evolves over time. Some genes, such as *Runx* are activated in founder cells and continue to be expressed during budlet development (Fig.5A) Others, like the transcription factors *Boschl.Bs8.g11353-SPP1* or the collagen *Boschl.Bs16.g20741-COL2A1*, stop being expressed in the early budlet and remain expressed only in peribranchial cells (Supp. Fig. 18), highlighting a transcriptomic shift from peribranchial identity to a budding specific program.

We then computed CytoTRACE scores to estimate the differentiation state of the budlet subset over successive stages. Among this subset, this budlet cluster exhibits a clear pattern with an increasing score from D to B1 followed by a decreasing score (Supp. Fig. 16C). This contrasts to the other clusters, which do not exhibit such a pattern and generally have lower scores. These results are consistent with a less differentiated, more progenitor-like state of the early budlet cells and a possible transition from peribranchial epithelium state to bud via cell dedifferentiation. (Fig.5D, Supp. Fig. 16C).

## Discussion

The multi-stage dataset presented here provides an unprecedented view of agametic reproduction in tunicates offering an important resource for understanding the cellular and molecular mechanisms underlying peribranchial budding. The dataset captured a wide diversity of cell types associated with budding and budlet formation, with clear annotation of major clusters supported by robust gene expression profiles. This not only validates previously described expression patterns but also enables high-resolution subclustering of complex tissues and allows to revisit developmental hypotheses with broader transcriptomic and genomic context.

One notable example is the endostyle, where transcriptomic zoning allows subregional identification, or revealing the nature of under-described anatomical structures like the peduncular septum, which likely derives from the peribranchial epithelium and remains connected to the budlet beyond stage B2 (double-vesicle). Similarly, developmental trajectories were observed across multiple lineages, including muscle and nervous systems, where gene expression gradients reflect known stage-specific differentiation events.

Despite the strengths of the dataset, some limitations must be acknowledged. For instance, it is important to note that some stages are less comprehensively represented than others. In particular, fewer cells were collected and sequenced during stages C1 and C2, periods marked by key events in tissue differentiation. Also, although the primary bud at stage C2 is nearing functional maturity, certain cell types or states specific to fully developed adults or to unrepresented vascular regions may be underrepresented or absent from our dataset. Additionally, the FACS gating strategy, which excluded haploid and potentially polyploid cells, resulted in the absence of mature gametes, limiting insights into the full germline spectrum.

In exploring the origin of cells contributing to budding, we focused on identifying candidate progenitors based on proliferation signatures, expression of germline multipotency program (GMP) genes, and homologs of Yamanaka factors (60). While tunicate budding has been linked to contributions from both mesenchymal and epithelial cells (*e.g.* (25)), our data provides no clear molecular evidence for a multipotent progenitor population. GMP marker expression was largely restricted to the germline cluster, and cells co-expressing *SoxB1* and *Myc* were limited to tissues such as the endostyle and stigmata, without forming a distinct pluripotent population. We also started exploring the role of the endostyle, previously proposed as a stem cell niche (26), and found expression of several *Wnt* genes, which is not sufficient to conclude with certainty whether it promotes a niche environment.

Given that in different colonial tunicates the putative stem cells involved in budding have been proposed to be circulating (2, 25, 27, 29, 72), we examined in greater detail the populations of circulating cells. In *Botryllus* specifically, grafting experiments demonstrated that rare, potentially circulating cells can give rise to either somatic or germline tissues, but not both (25). By comparing our budding dataset with a reference dataset of circulating cells (sampled at stages A1 and B2), we identified the clusters corresponding to cell types present in circulation. As expected, both germline and somatic cell populations were represented. Notably, the germline cluster appeared transcriptionally distinct and distant from the somatic clusters, consistent with the well-documented separation between somatic and germline lineages in tunicates(73) and previous grafting experiments (25).

Interestingly, a substantial number of the circulating cell clusters form a transcriptionally interconnected group, suggestive of ongoing differentiation trajectories. Within this group, we identified two clusters likely corresponding to the previously described hemoblasts, undifferentiated blood cells thought to harbor stem-like potential (67).

The diversity of cell populations revealed by clustering exceeds the eight morphotypes historically described based on morphology (74). Moreover, genes previously associated with a specific hemocyte morphotype, such as the phenoloxidase *Ppo1-like* and *Botryllin* for morula cells, are expressed in different clusters, indicating a previously unappreciated molecular heterogeneity in this morphotype. This higher resolution is more consistent with the populations identified by Rosental and colleagues using FACS-based profiling, and future efforts could aim to align those sorted populations with our transcriptomic clusters (63).

Despite the identification of putative hemoblast clusters, PAGA trajectory analysis did not reveal strong developmental links between these clusters and the main tissues of the bud, though lowering the threshold suggested possible connections between cluster 23 (resident hemoblast-like cells) and cluster 12, and connections between an unidentified hemocyte cluster (cluster 38), and cluster 20 (late branchial/peribanchial and stalk septum cells) or cluster 13 (*Tgf-ꞵ-f*⁺), the significance of these connections and whether they reflect an actual biological link need further analysis.

Altogether, these observations indeed support the presence of circulating stem-like cells in *B. schlosseri*. However, our dataset does not provide evidence that these cells contribute broadly to somatic tissues of the bud beyond the hematopoietic lineage or the somatic and germline compartments of the gonads.

A particular intriguing observation is the transcriptional diversity within the *Tgf-ꞵ-f*⁺ cell population, which appears to comprise at least three transcriptionally distinct subtypes. These may correspond to different functional identities, potentially including support cells, somatic components of the testis, or follicular cells (32). A less conservative PAGA trajectory analysis revealed proximity of these clusters to early neuron and heart lineages (Supp. Fig. 6E), suggesting possible developmental or functional links. This raises the hypothesis that certain *Tgf-ꞵ-f*⁺ populations may contribute to, or interact with, specific organ lineages during organogenetic phases of budding, rather than representing a uniform cell type.

To identify the founder cells of the bud, we searched for a transcriptionally distinct population within the early peribranchial epithelium. We identified a small group of cells characterized by a previously described specific combination of gene expression features (5, 12, 23, 32). CytoTRACE analysis inferred an increase in developmental potency from these founder cells from budlet onset (stage D) toward the budlet stage B2. Since CytoTRACE scores are partly based on the number of genes expressed, often higher in less differentiated cells with more open chromatin (35), this trend suggests a progressive gain of potency during the early stages of budding. While transcriptional data alone cannot definitively determine cell potency, these findings imply that the founder cells are not intrinsically multipotent but may acquire such potential through reprogramming mechanisms during early bud development. This aligns with classical features of dedifferentiation, which typically involve changes in gene expression programs, protein composition, cell morphology, and potency (75).

Our data further reveal a marked switch in transcriptomic identity in these founder cells. Genes typically expressed in the peribranchial epithelium, such as the transcription factor *Pax2/5/8a*, the PHF zinc finger protein *Boschl.Bs8.g11353-SPP1*, and the collagen *Boschl.Bs16.g20741-COL2A1*, are downregulated. Concurrently, other genes become upregulated, including *Runx* and, slightly later, *Nk4*. Within the peribranchial epithelium at stages C2 and D, *SUMO2* (*Boschl.Bs3.g4309-SUMO2.3*) is expressed exclusively in the founder population. SUMOylation is a post-translational modification involved in diverse cellular processes (76), and has been shown to play a key role in stabilizing both stemness and differentiated cell states (77, 78).

Additional genes involved in post-transcriptional and proteome regulation are also enriched in the founder cells and early budlets, including the splicing regulator *Mbnl* and the E3-ubiquitin ligase *Boschl.Bs7.g9551*, suggesting active remodeling of the proteome and potential regulation of cell state transitions.

Several signaling pathways have been implicated in budding in styelid ascidians such as *B. schlosseri*, although functional studies often yield complex phenotypes that are challenging to interpret. While we cannot directly infer pathway activation from our data, we examined the expression of potential receptors that may be involved in bud induction. Notably, a *Gnrhr*-like gene is transiently expressed in founder cells and early budlets, as well as in the gonads, nervous system, and other cell populations. In vertebrates, gonadotropin-releasing hormones (GNRHs) and their receptors (GNRHRs) regulate the hypothalamus–pituitary–gonadal axis, particularly in response to the organism’s metabolic state (79). In invertebrates, which lack this axis and gonadotropins, GNRHs are thought to act directly as neuropeptides modulating both reproductive and non-reproductive functions (80). In *Ciona*, GNRH signaling has been implicated in both reproduction and metamorphosis. In *Botryllus*, the number of buds and the extent of gonad development are known to vary with environmental conditions, a well as with others factors such as age(7). These observations raise the possibility that a GNRH-like signaling axis may play a dual role in coordinating budding and reproductive development in response to the colony’s metabolic state.

## Conclusions

Our multi-stage single-cell atlas of *Botryllus schlosseri* agametic development provides the first transcriptomic roadmap of peribranchial budding at cellular resolution. Our data identify a distinct founder population emerging from the peribranchial epithelium, supporting a model of budding triggered by context-dependent dedifferentiation and remodeling of committed somatic tissues rather than a fixed pool of multi-/pluripotent stem cells. While we identified circulating hemoblast-like populations with potential stem-like features, our analysis did not reveal clear broad contributions of these cells to bud onset beyond the hematopoietic or gonadal lineages. Instead, our data suggest that the peribranchial epithelium itself serves as a source of developmental plasticity, acquiring gene expression signatures indicative of increased potency through successive stages, at least in the budlet and its early morphogenesis. Moreover, the unexpected molecular diversity within traditionally defined cell types, such as hemocytes and Tgf-ꞵ-f⁺ populations, highlights the complexity of cellular specialization during asexual development. By integrating trajectory analyses, founder cell signatures, and pathway expression profiles, we also propose candidate regulatory mechanisms, including GNRH-like signaling, that may couple environmental and metabolic cues to the initiation of budding. Altogether, this work expands our understanding of non-embryonic development in chordates and establishes a rich resource for future studies of regeneration, stemness, and plasticity in a vertebrate sister group model.

---

**Supplementary Figure 1.**
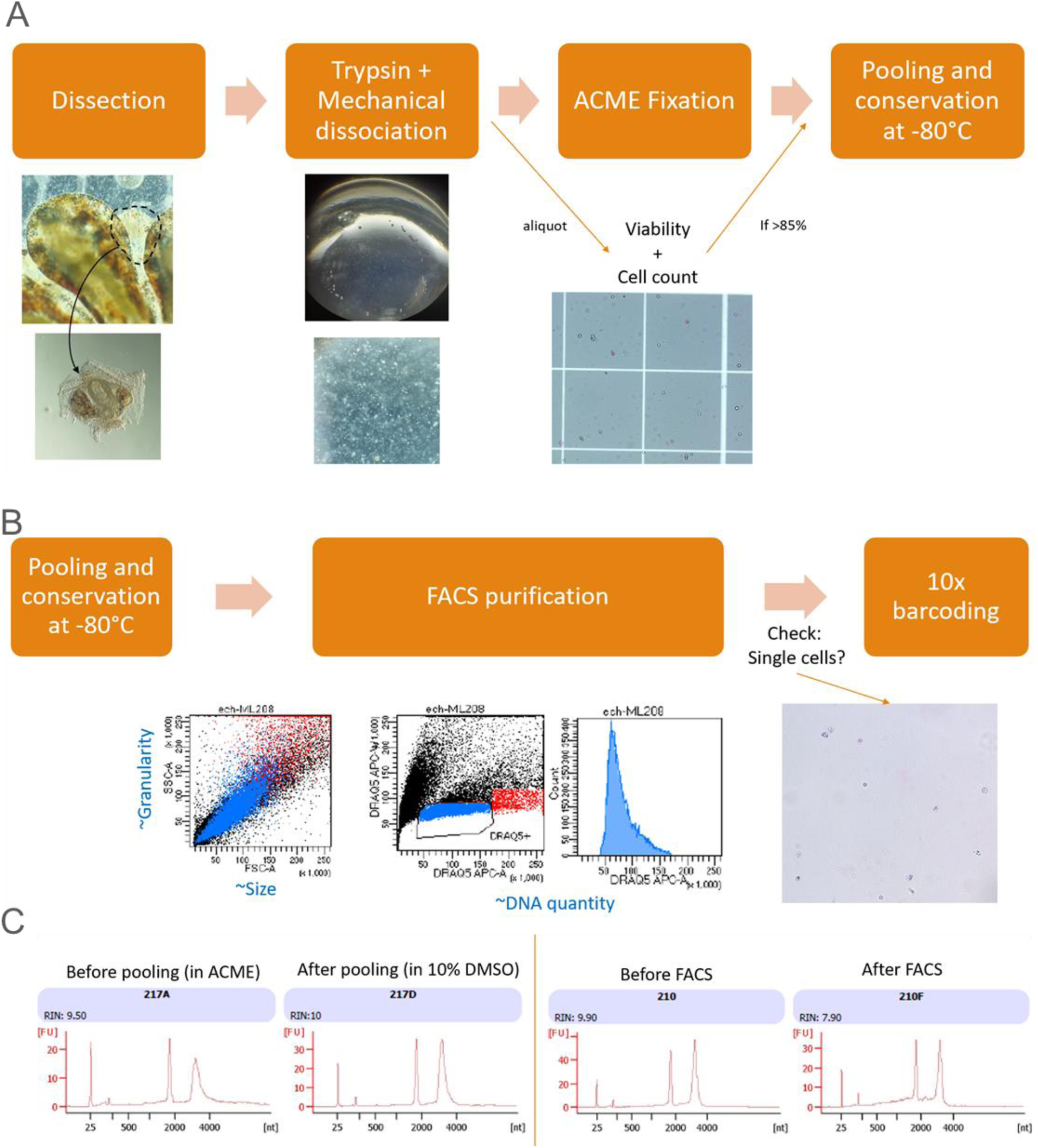
Preparation of the samples for the full budding scRNAseq dataset. (A) Main steps of the sample preparation shown for one sample, from dissection to preservation at −80°C. (B) FACS gating based on DNA content (stained with Draq5) to remove debris and cell aggregates. (C) Bio-Analyzer chromatograms representatives of the different samples dissociated and tested for RNA quality, both before and after cryopreservation, and before and after FACS. Raw data and viability measurements for each sample are available upon request.

**Supplementary Figure 2.**
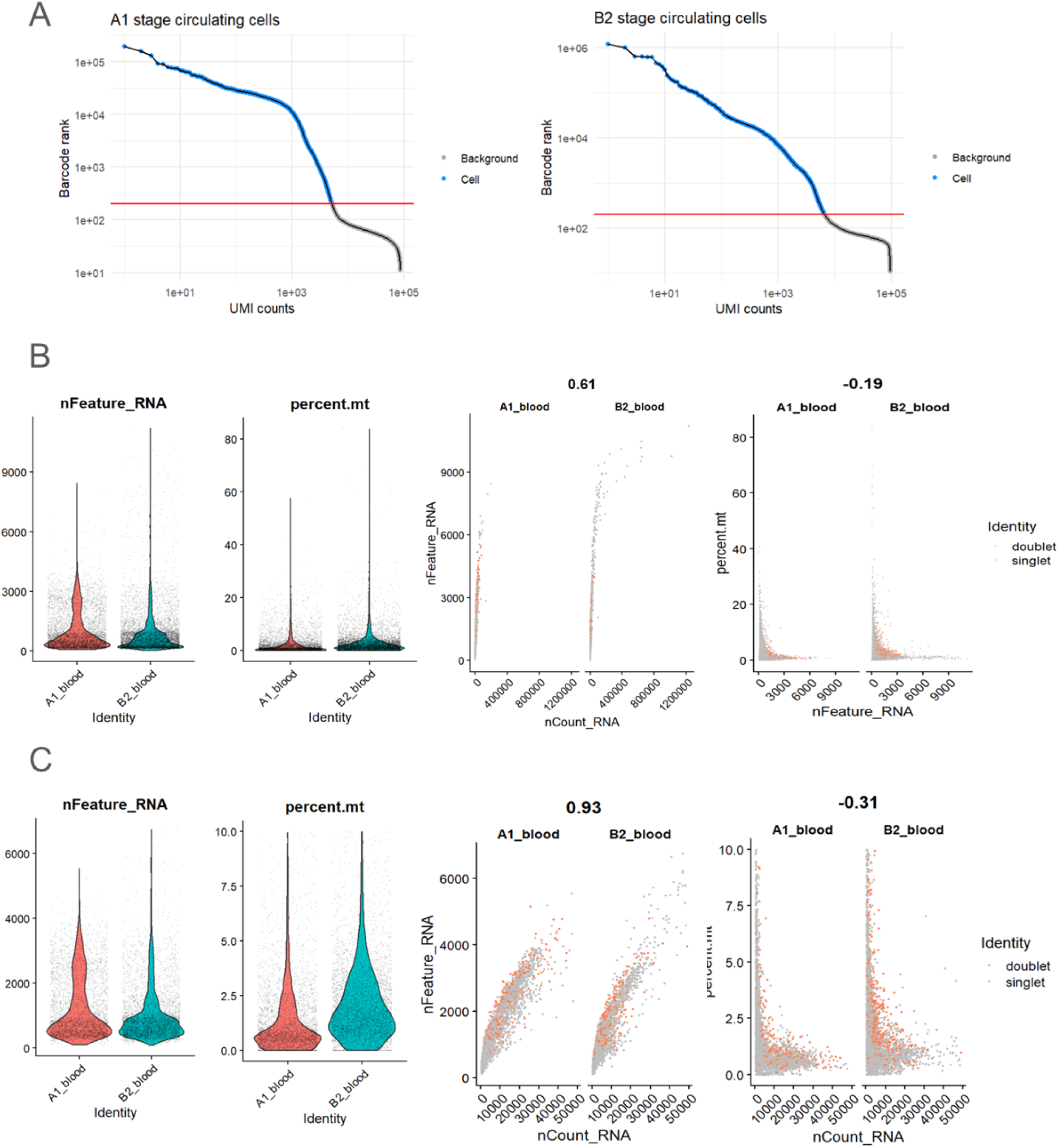
Distribution of the quality metrics in each circulating cell sample pre- and post-filtering. **(A)** Barcode rank plots of the two circulating cell samples showing the threshold to filter inferred empty droplets. Red line = 200 UMIs threshold. **(B-C)** Distribution of quality control metrics **(B)** pre- and **(C)** post-filtration. Cells were kept if the number of detected UMIs (nCount_RNA) was between 500 and 50000, and the percentage of UMIs from mitochondrial genes (percent.mt) was under 10. We decided against filtering out the doublets in these samples because contrary to expectations, the “doublets” detected by scDblFinder are evenly distributed, and not concentrated toward high UMI counts/high gene counts. In addition, due to the nature of the samples (circulating cells), only rare multiplets are expected

**Supplementary Figure 3.**
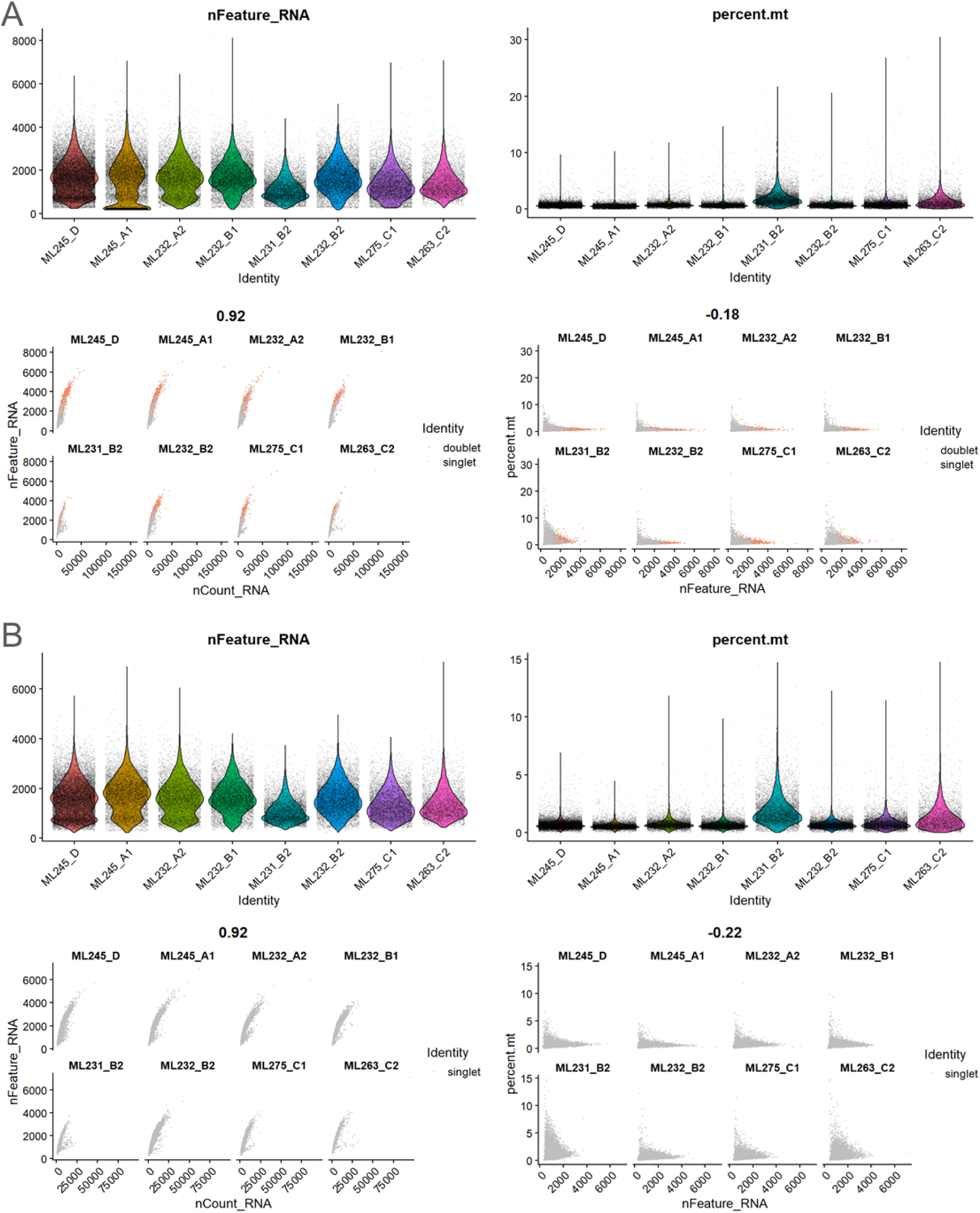
Distribution of the quality metrics in each budding sample pre- and post-filtering. (A-B) Distribution of quality control metrics **(A)** pre- and **(B)** post-filtration. Cells were kept if the number of detected UMI (nCount_RNA) was higher than 750, the percentage of UMI from mitochondrial genes (percent.mt) under 15%, and either the gene count (nFeature_RNA) over 400 or the percent.mt under 5.

**Supplementary Figure 4.**
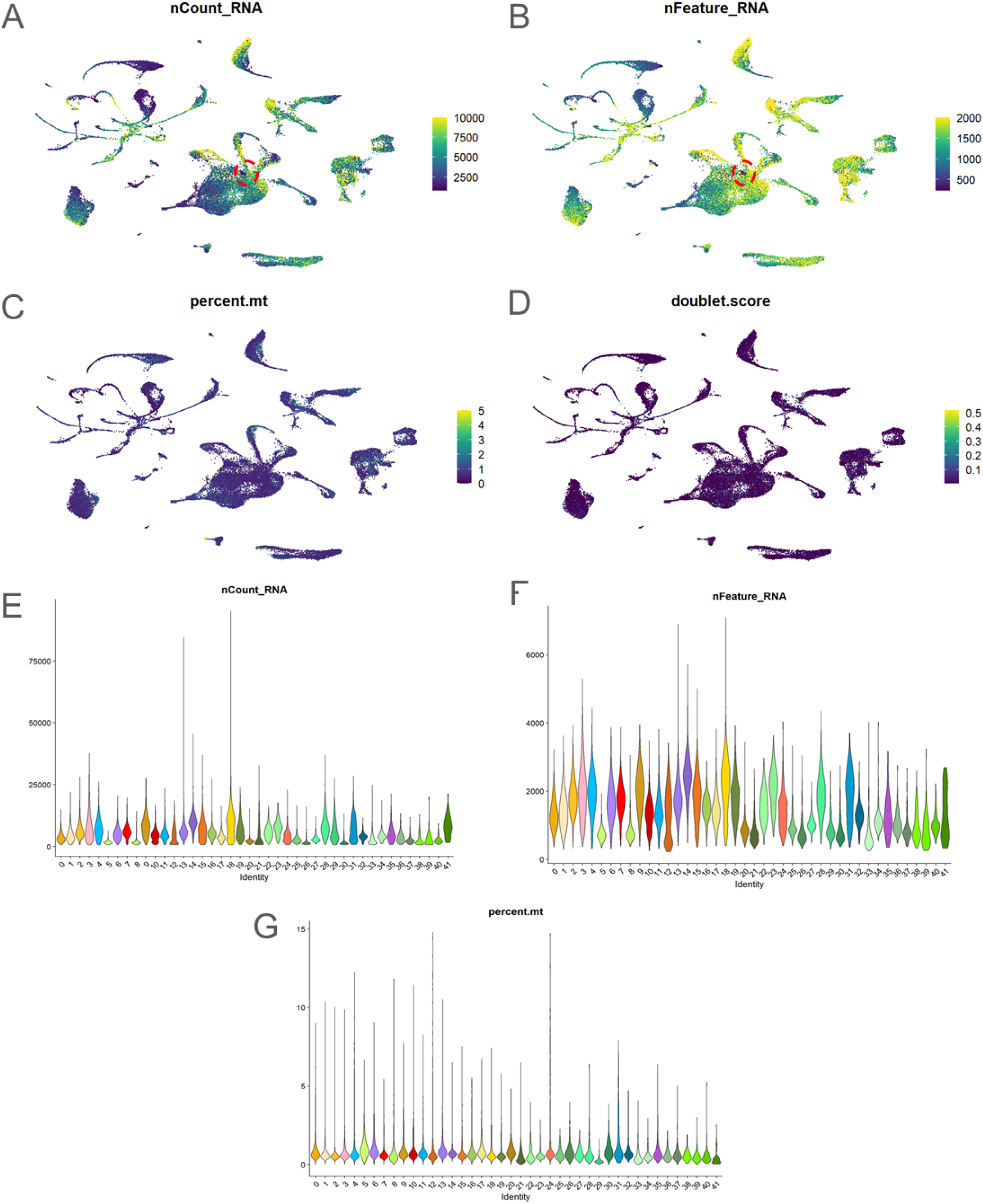
Quality metrics of the full budding dataset. (A-D) Distribution across the UMAP and (E-G) violin plots of quality metrics. Most clusters show a good distribution of the quality metrics. A portion of cluster 12 might be cells of lower quality, as it has both very low UMI number and gene counts (dashed red circle in A, B). (A, E) Total number of RNA molecules (UMIs) per cell (the UMAP color max was set at 10,000 for readability). (B, F) Genes counts per cell (the UMAP representation max was set at 2,000 for readability). (C, G) Percentage of mitochondrial RNA molecules per cell (the UMAP color max was set at 5% for readability). (D) Doublet score assigned by *scDblFinder*. All cells correspond to singlets. No high doublet score cluster is visible, suggesting good filtering.

**Supplementary Figure 5.**
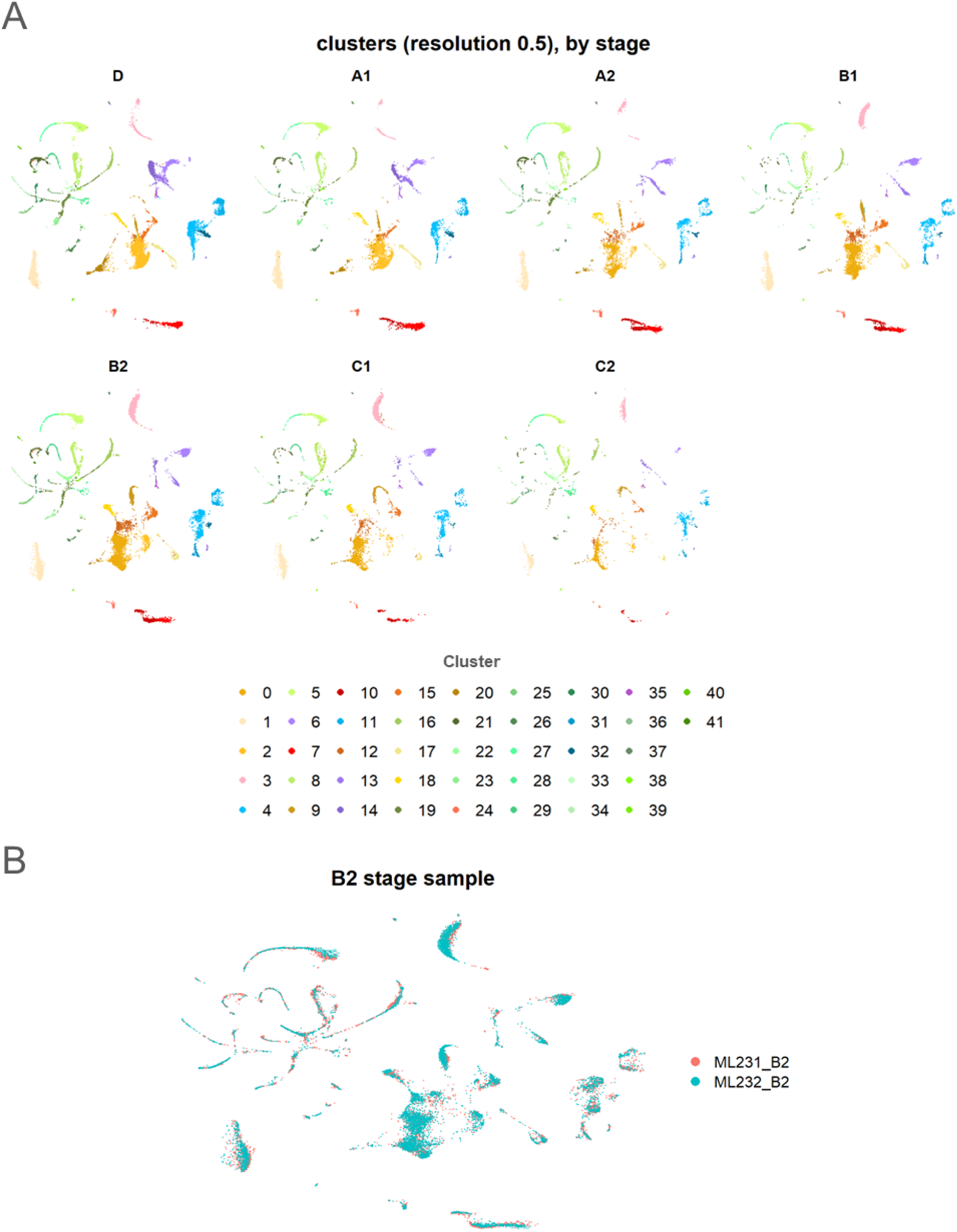
Distribution of the stages and the two B2 stage replicates across the dataset. (A) UMAP projection of the full budding dataset split by stage. (B) UMAP projection with only the two B2 samples replicates. The B2 samples (prepared and sequenced at different times on the same clonal colony) are superposed almost perfectly on the UMAP, suggesting a low batch effect in our dataset.

**Supplementary Figure 6.**
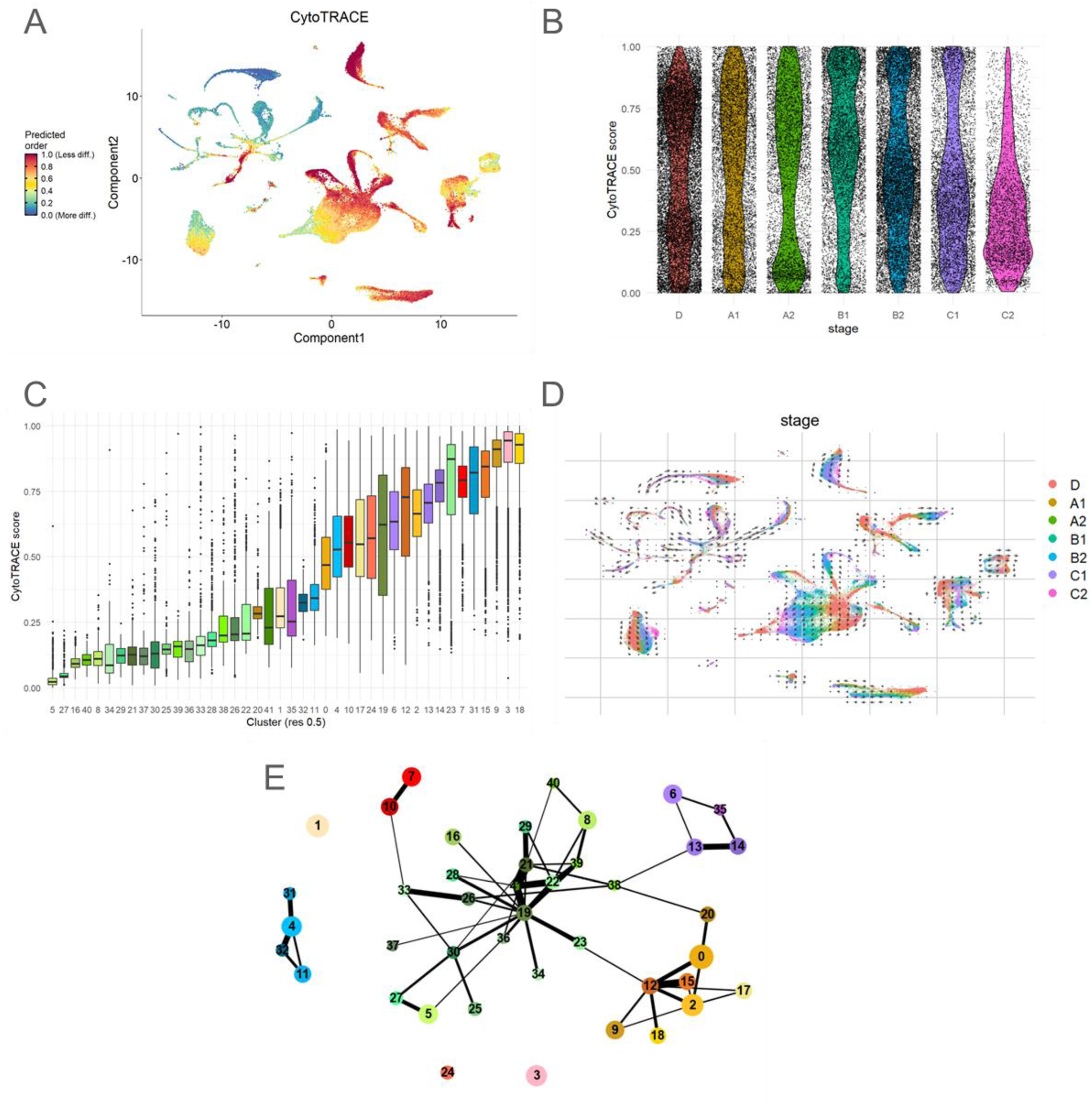
CytoTRACE, RNA velocity and PAGA analyses of the full budding dataset. (A, **C)** CytoTRACE score, **(A)** Distribution of the score over the full budding dataset, **(B)** Score distribution by stage, **(C)** Score distribution by cluster (resolution 0.5). **(D)** RNA velocity projected on the UMAP, with colors representing stage. **(E)** Abstracted graph (PAGA) of the full budding dataset, with PAGA threshold=0.1, showing some low strength connections between the “hemocytes” module (green) and the other modules.

**Supplementary Figure 7.**
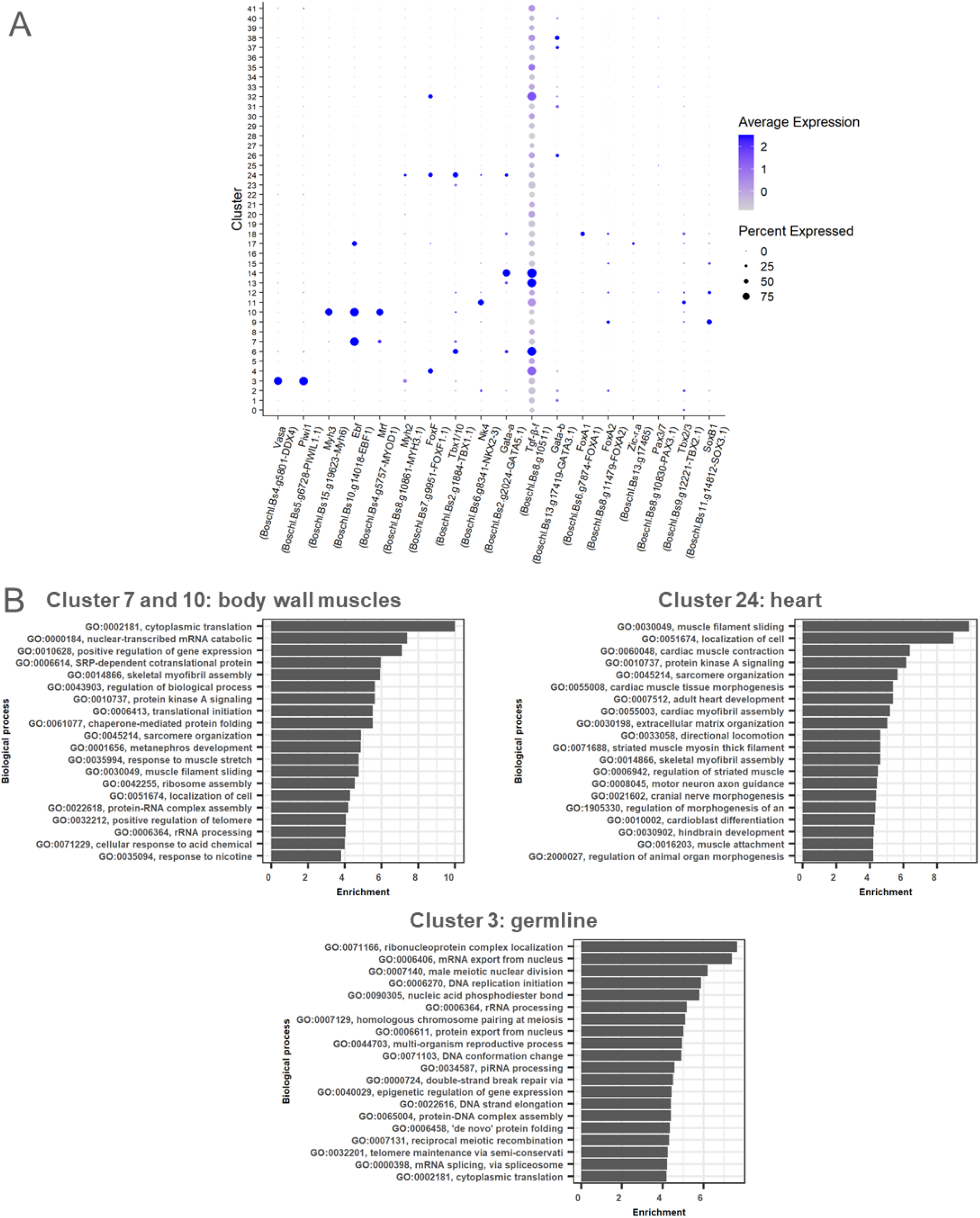
Annotation of clusters identity from published expression patterns and GO term enrichment. **(A)** Scaled expression of genes with described expression patterns in *B. schlosseri* and other tunicates. The genes correspond to the ones presented in Supplementary Table 3, except the cellulose synthase gene *CesA*, which was obtained by protein BLAST from *Ciona intestinalis CesA* (NP_001041448.1). Expression of the gene is scaled. **(B)** GO term enrichment analysis of the body wall muscles, heart, and germline clusters.

**Supplementary Figure 8.**
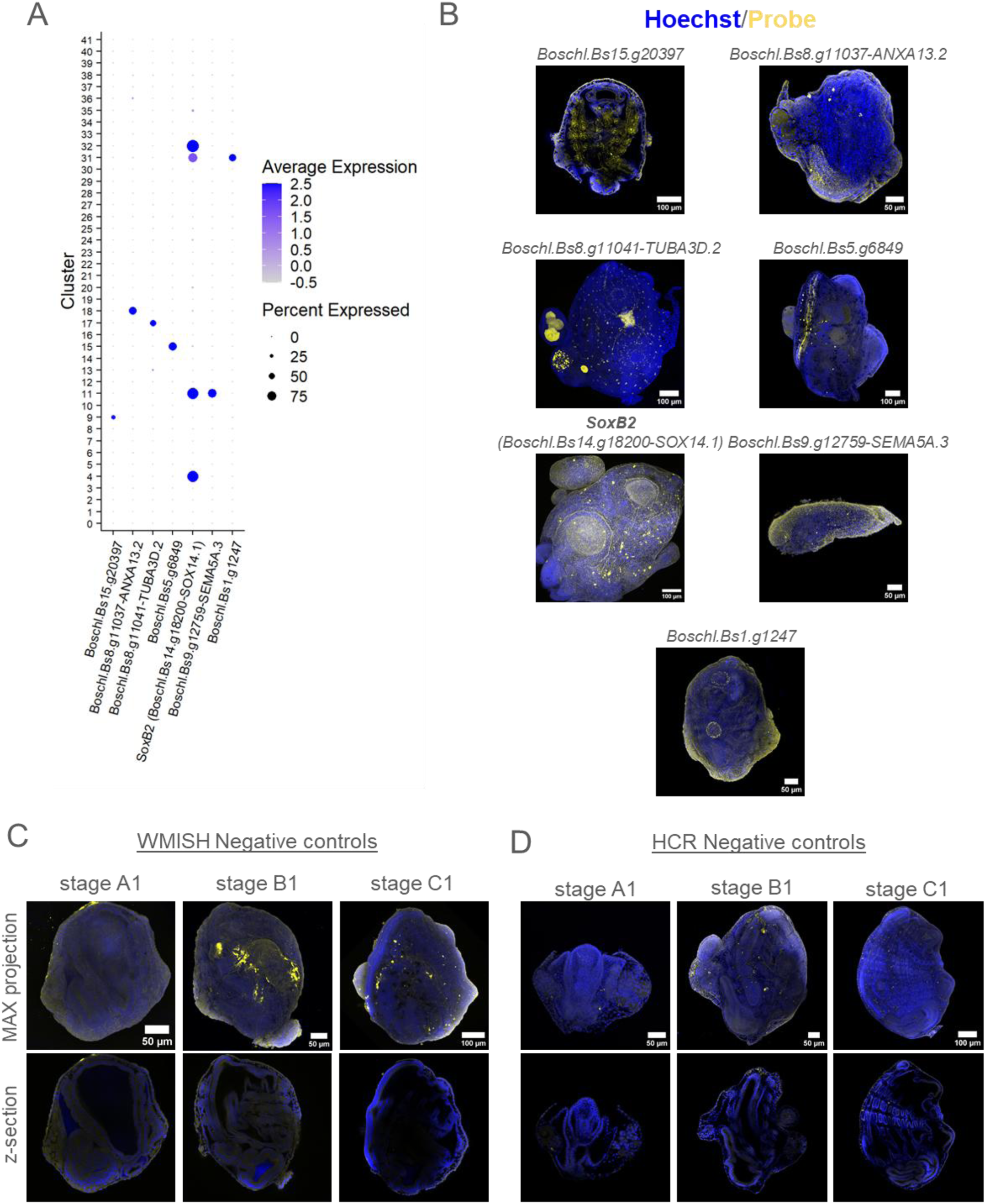
Additional gene expression patterns to annotate the clusters, and negative controls. (A) Scaled expression of additional marker genes specific to a tissue type, in the different clusters of the full budding dataset (resolution 0.5), and (B) corresponding expression patterns observed by WMISH in primary buds or in ampullae. (C-D) Negative controls for (C) WMISH and (D) HCR. Some mesenchymal cells with specific morphology are fluorescent in the FITC (C) or Alexa-647 (D) emission window.

**Supplementary Figure 9.**
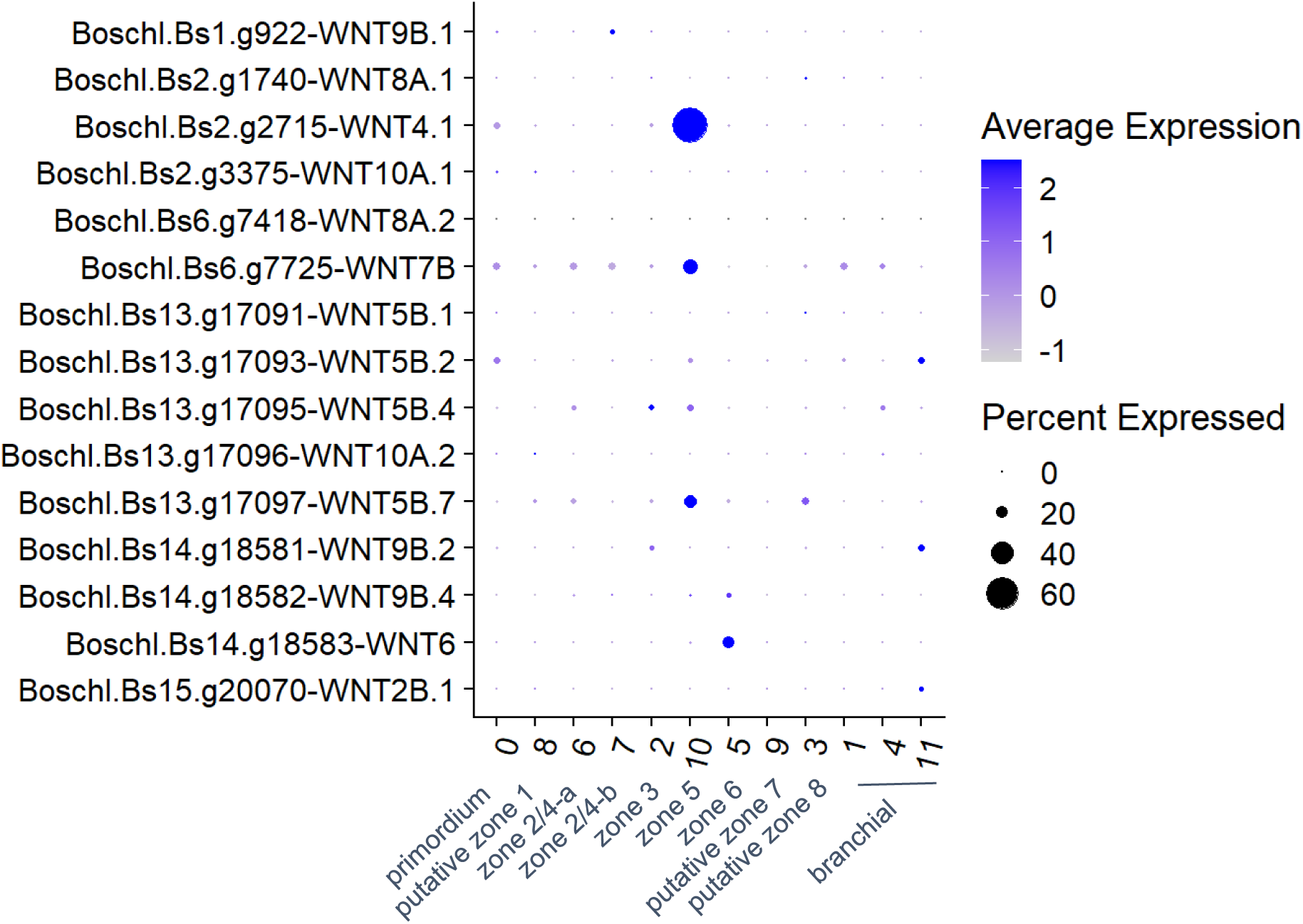
Scaled expression of WNT genes in the endostyle subset. The presented Wnt genes were selected from the functional annotation. Clustering correspond to the one presented in Fig. 3.

**Supplementary Figure 10.**
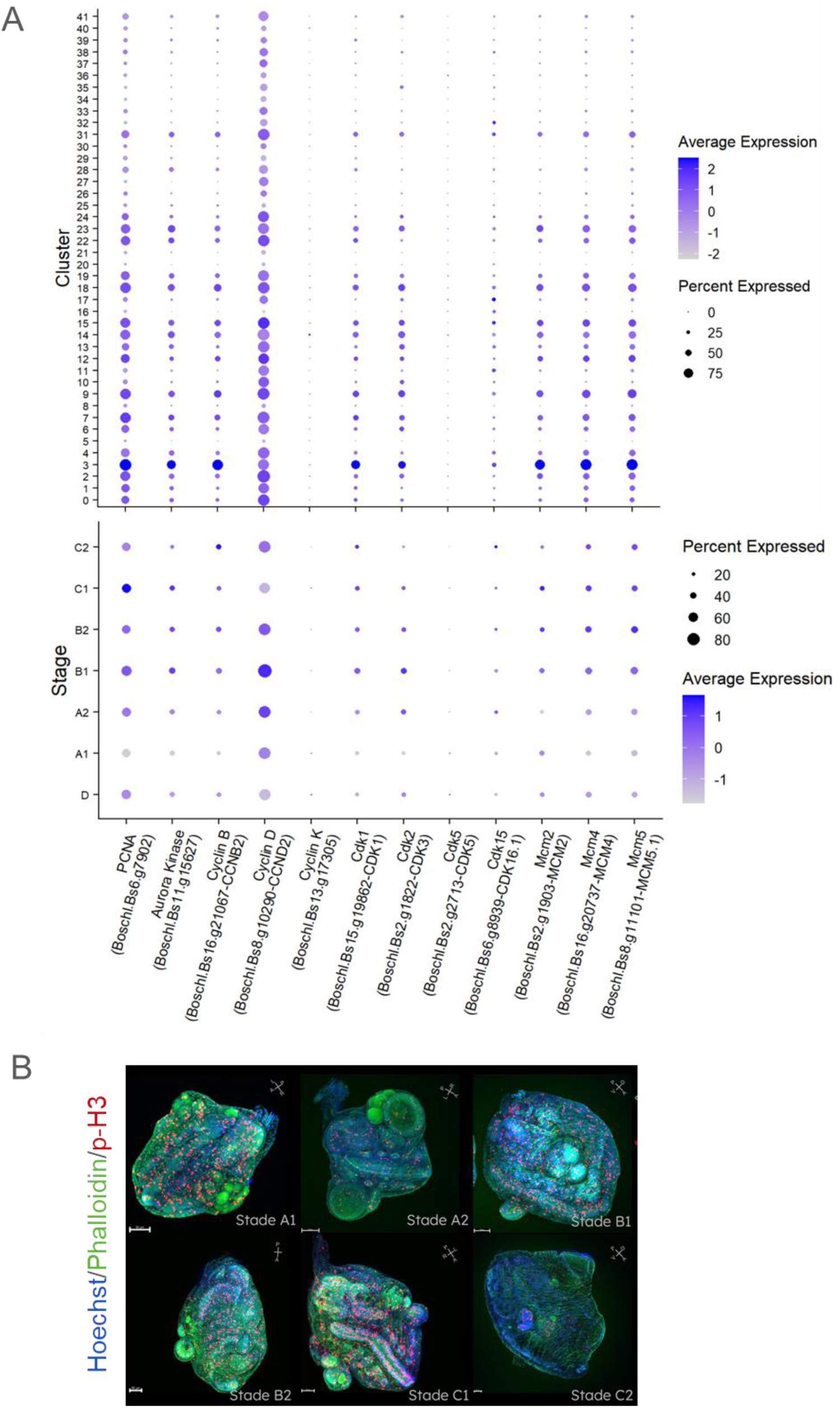
Proliferative dynamic in the budding dataset. (A) Scaled expression of genes associated with proliferation, by cluster (top) and by stage (bottom). (B) Confocal pictures of buds and budlets stained with an antibody against phospho-histone H3, phalloidin and Hoechst. Orientation is shown on the top right, scale bar is 50µm.

**Supplementary Figure 11.**
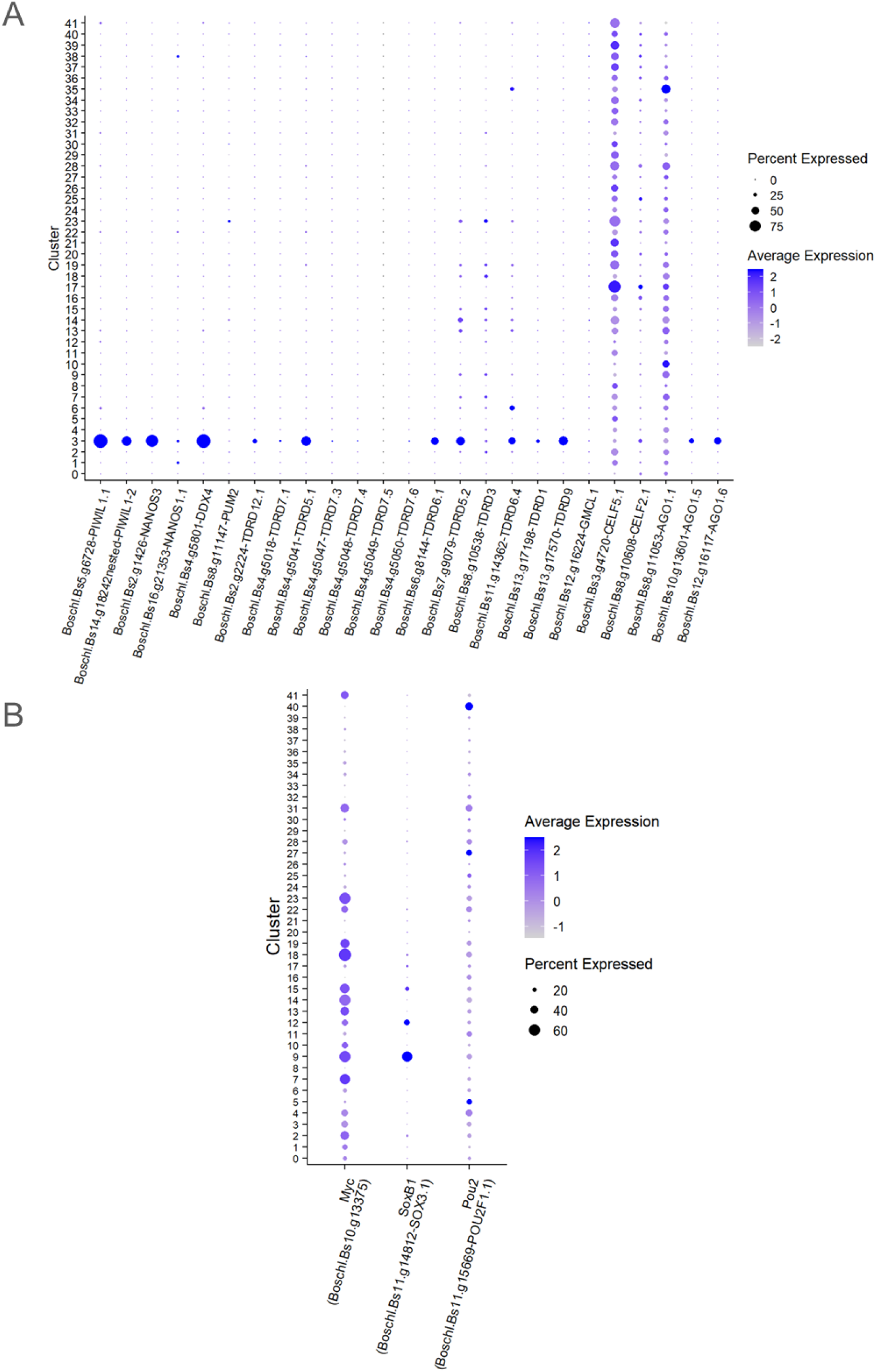
Expression of gene sets associated with adult stem cells. (A-B) Scaled expression (A) of germline multipotency program genes and (B) of Yamaka factors-related genes in the clusters of the full budding dataset (resolution 0.5).

**Supplementary Figure 12.**
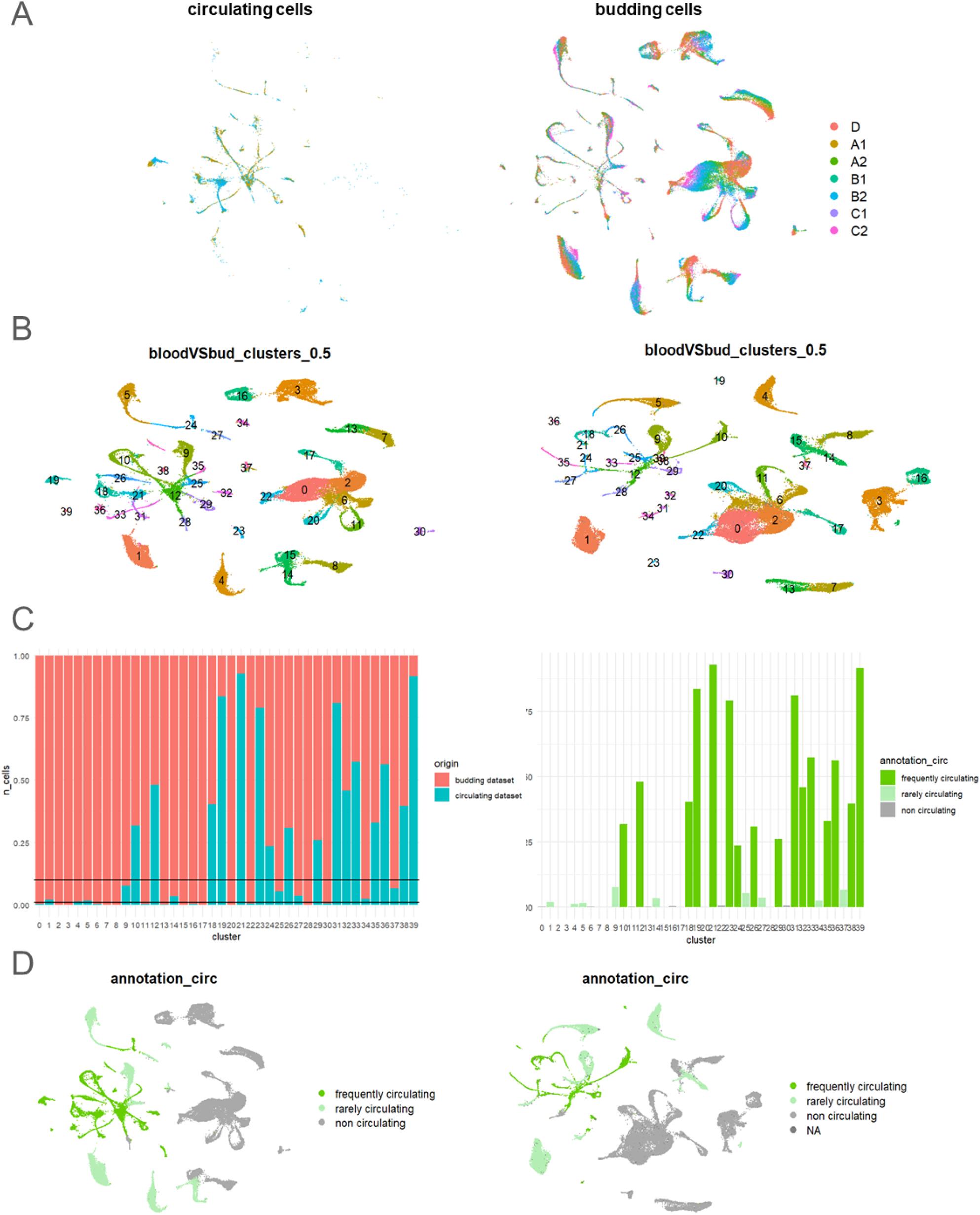
Comparison of budding and circulating cells datasets. (A) UMAP projection of the joint dataset, colored by stage and split by dataset of origin. (B) UMAP projection of the joint dataset (left) and the budding dataset (right), showing the correspondence between cluster numbers (resolution 0.5). (C) Proportions of cells coming for each dataset in each cluster (left), and corresponding annotation on circulating status (right). (D) UMAP projection of the joint dataset (left) and the budding dataset (right), colored by circulating status.

**Supplementary Figure 13.**
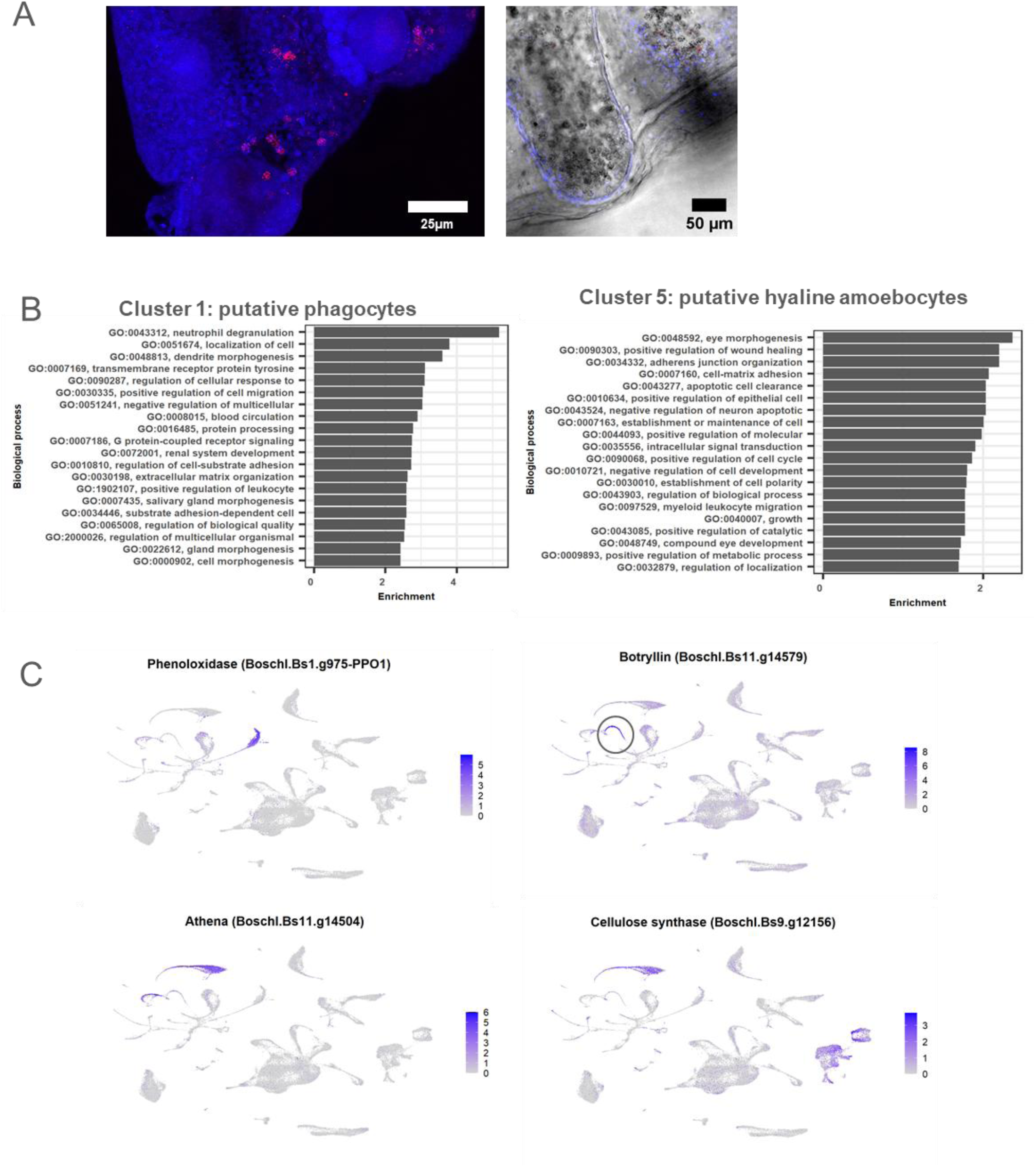
Additional analysis of circulating cells clusters. (A) Negative controls of HCR showing autofluorescent morula cells in a primary bud, and no autofluorescent cells in the tunic. (B) 20 most significantly enriched Gene Ontology terms in various hemocyte clusters. (C) Expression plots of various genes known to be expressed in circulating cells.

**Supplementary Figure 14.**
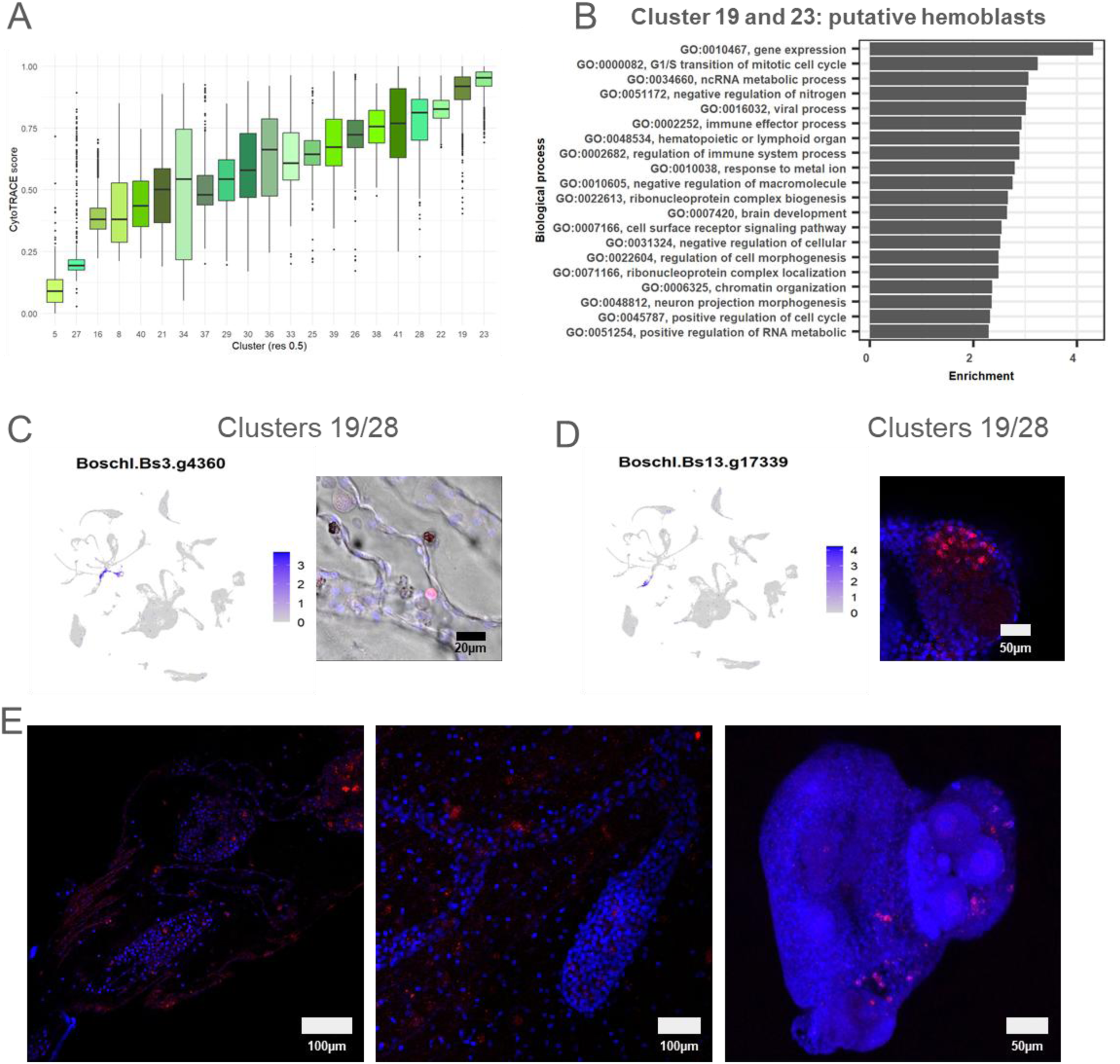
Additional analysis of the putative hemoblasts. **(A)** CytoTRACE scores computed on the hemocyte subset, per cluster. **(B)** 20 most significantly enriched Gene Ontology terms in the clusters 19 and 23 compared with the rest of the dataset. **(C-D)** Expression plots and HCR of marker genes of putative hemoblasts (cluster 19) and another hemocyte cluster (cluster 28) **(C)**, or resident hemoblast-like only **(D)**. **(E)** Negative controls of HCR showing autofluorescent morula cells. For HCR pictures: the probe signal is in red, Hoechst staining in blue, and the transPMT in grey.

**Supplementary Figure 15.**
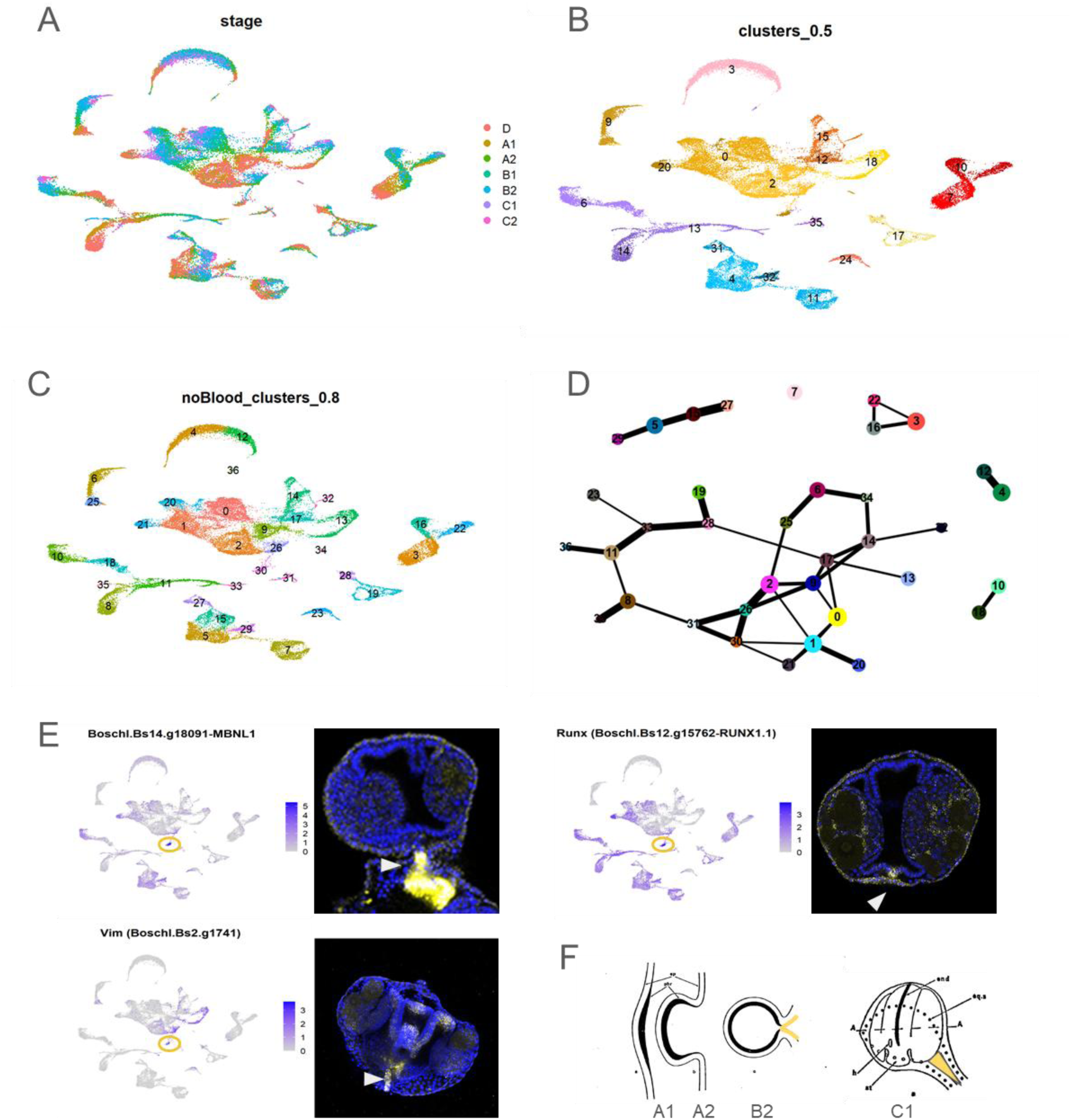
Additional analyses to search for the budding founder cells. (A-C) UMAP projection of the subset without hemocytes and putative phagocytes, showing (A) stages, (B) cluster numbers from the full budding dataset and (C) cluster numbers of this subset subclustering at resolution 0.8. (D) Corresponding abstracted graph (PAGA). (E) Expression plots and HCR of genes expressed in the budlet stalk septum; the stalk septum cluster (cluster 30) is circled in yellow. (F) Schematics of the formation of the budlet stalk (yellow), modified from (71).

**Supplementary Figure 16.**
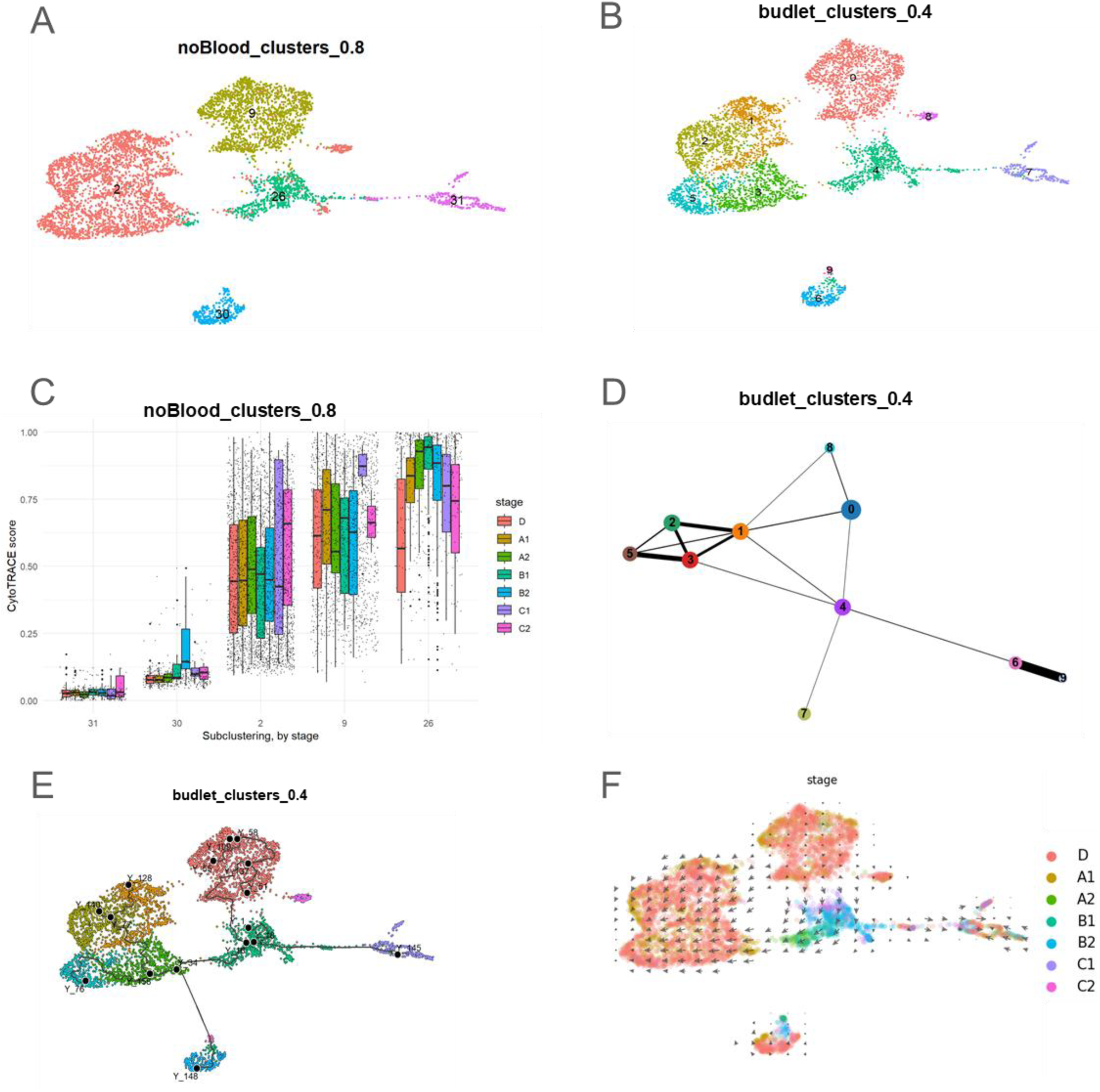
Budlet founder cells are a subset of the peribranchial population. (A) UMAP projection subset containing early branchial (9) and peribranchial (2), budlet (26), budlet stalk septum (30) and Bhf+ (31) cells. The cluster numbers correspond to the ones in Sup Fig 16B. (B) Subclustering of the “budlet” subset. (C) CytoTRACE score of the different clusters displayed in (A), by stage. (D) Abstracted graph (PAGA) of the clusters of the “budlet” subset (see B). (E) Monocle3 inferred trajectories for the “budlet” subset. (F) Projected RNA velocities vectors for the “budlet” subset.

**Supplementary Figure 17.**
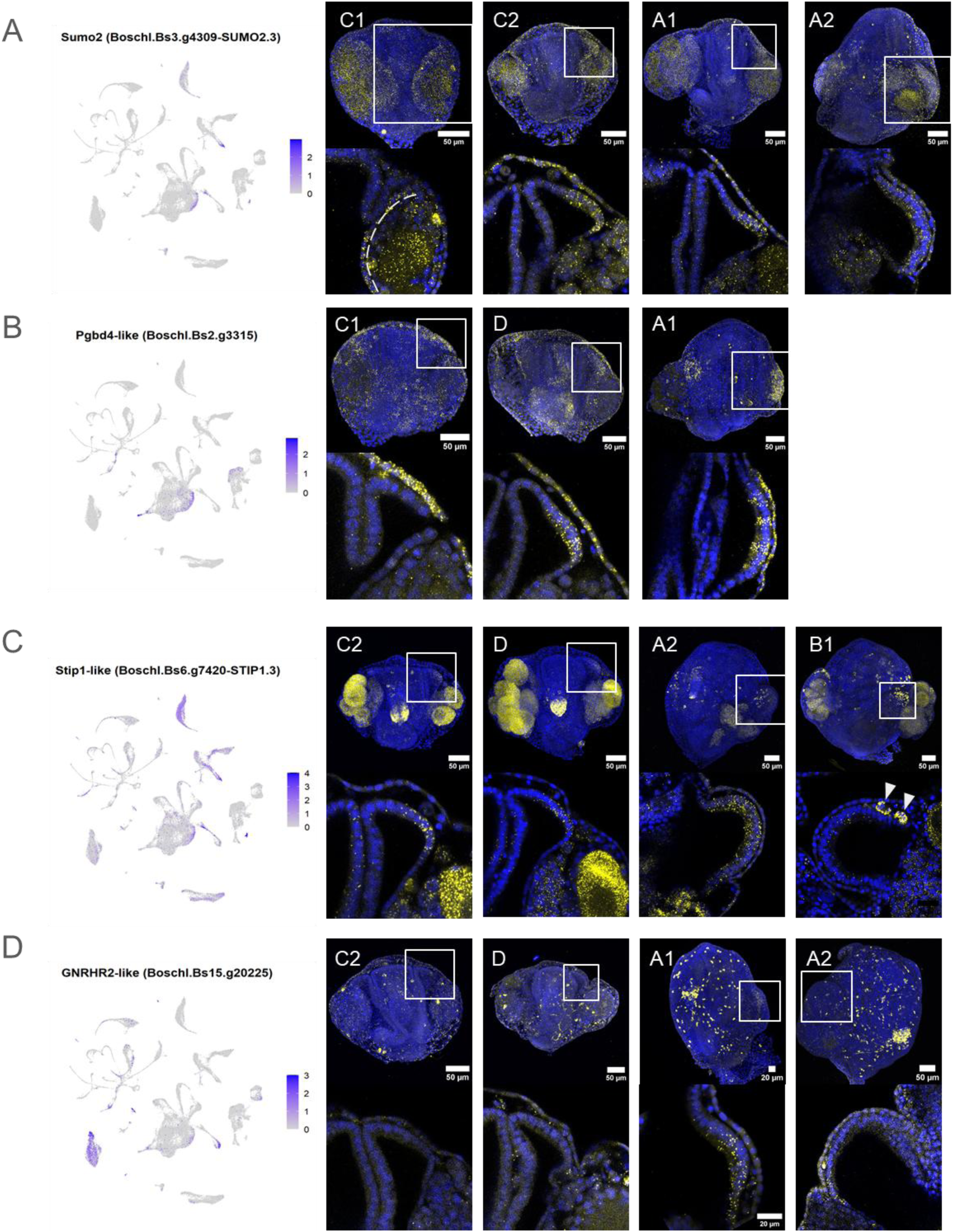
Expression patterns of the budding founder cells markers over time. (A-D) Expression plots and confocal images of HCR on genes enriched in the budding founder cells. The top rows are maximum projections of the whole z-stack, The bottom rows correspond to a confocal section of the budlet area squared in the top row image. Stage is on the top left. (A) *Sumo2-like* (Boschl.Bs3.g4309-SUMO2.3) is expressed from stage C1, in the external side of the peribranchial primordia, and it continues to be expressed until stage A2. The limit between the peribranchial epithelium and the gonad is highlighted by dashed curve. *Sumo2-like* is also expressed in maturing oocytes, and in some Tgfβ-f+ cells in the gonads (B) *Pgbd4-like* (Boschl.Bs2.g3315) first expression in the peribranchial primordia has been detected at stage D. *Pgbd4-like* is also expressed in the budlet epidermis, in the stalk septum (not shown), and in some hemoblasts like circulating cells (not shown) (C) *Stip1-like* (Boschl.Bs6.g7420-STIP1.3) first expression in the peribranchial primordia has been detected at stage C2 at the anterior-external side of the right peribranchial epithelium. *Stip1-like* positive mesenchymal cells near the bud (white arrowheads) probably correspond to migrating germline or to Tgfβ-f+ cells, consistent with its high expression in cluster 3 and 13 (see Fig. 1). *Stip1-like* is also expressed in the maturing nervous system (D) *Gnrhr2-like* (Boschl.Bs15.g20225) is expressed at in the budlet and the peribranchial primordia just posterior to it from the early stage A1, and can still be detected at the posterior ventral side of the budlet at stage A2 (additional budlet primordium, developing later) the of the right peribranchial epithelium. It is also expressed in the nervous system as well as the cells of cluster 23, with a similar expression pattern as observed from genes of this cluster (see stage D image and Fig. 4K).

**Supplementary Figure 18.**
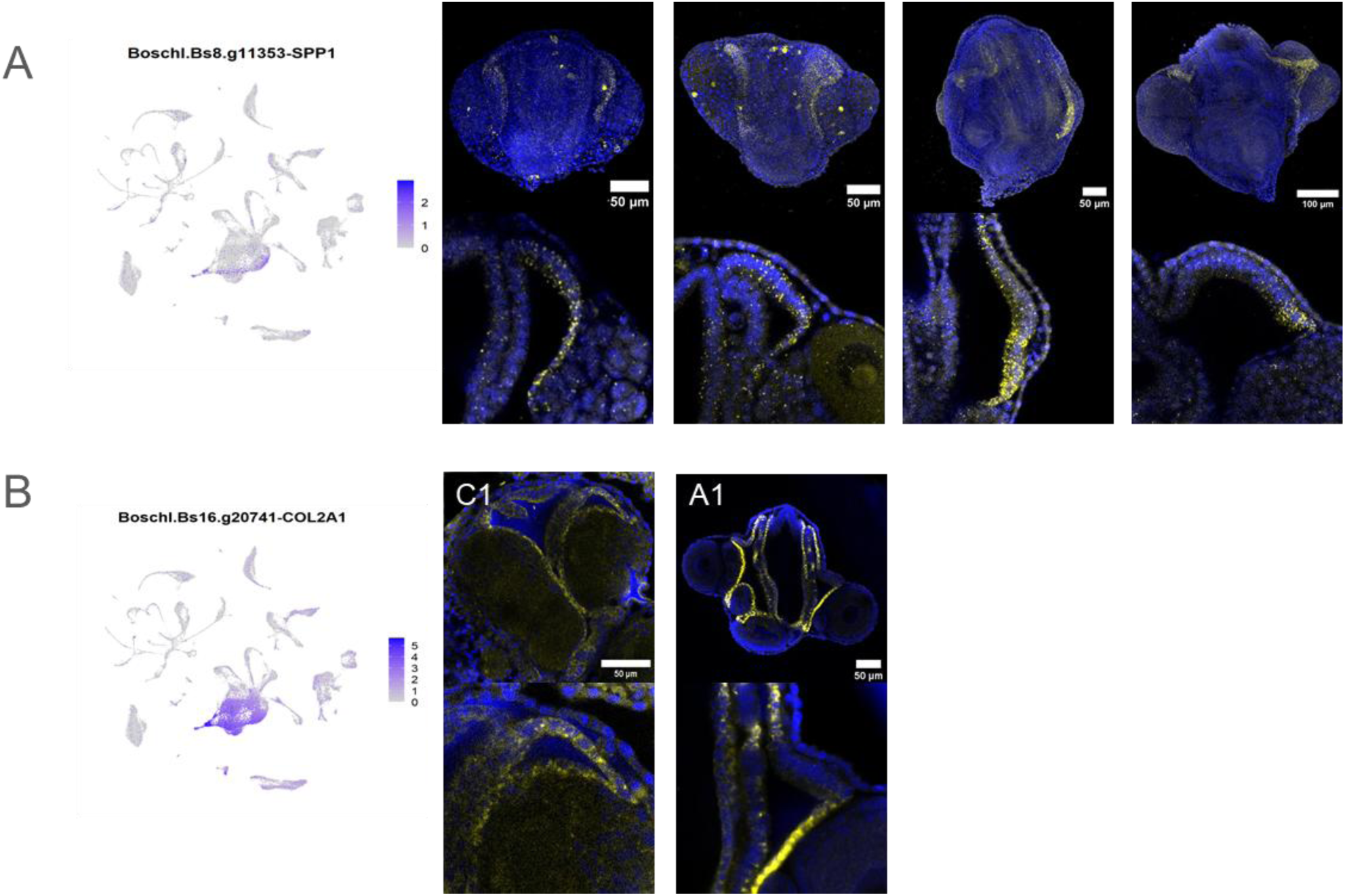
Expression pattern of peribranchial epithelium genes that stop being expressed in the budlet. (A-B) Expression plots and confocal images of (A) B*oschl.Bs8.g11353-SPP1* HCR and (B) the collagen *Boschl.Bs16.g20741-COL2A1* WMISH. Both genes are present in the budding founder cells, but depleted in the budlet compare to the peribranchial epithelium at later stages. The bottom rows correspond to a confocal section of the budlet area squared in the top row image. Stage is on the top left.

**Supplementary Table 1:** Summarized dissociation metrics for each sample in the budding dataset. The sample ID is the same as the one used in the Seurat object. Unless otherwise mentioned in the comment column, samples were obtained by pooling dissociation batches from the same day and colony. The number of singlets and viability were estimated on a 10µl aliquot of each dissociation batch (100-150µl total, 10-12 bud/budlets each) before fixation.

| Samples | Stage | Number of buds pooled | Estimated number of singlets before fixation | Average number of singlets/bud | Average viability | Pooled dissociations |
| --- | --- | --- | --- | --- | --- | --- |
| ML231-B2 | B2 | 53 | 905,400 | 17,083 | 93,8 |  |
| ML232-B2 | B2 | 65 | 1,234,240 | 18,988 | 93,6 |  |
| ML232-B1 | B1 | 83 | 509,600 | 6,140 | 96,0 | from 2 different days |
| ML232-A2 | A2 | 156 | 686,880 | 4,403 | 91,5 |  |
| ML245-A1 | A1 | 209 | 506,160 | 2,422 | 90,3 | from 2 different days |
| ML245-D | D | 337 | 694,000 | 2,059 | 89,38 | from 4 different days |
| ML263-C2 | C2 | 50 | 1,193,840 | 23,876 | 94,98 |  |
| ML275-C1 | C1 | 82 | 2,161,440 | 26,359 | 95,94 | from 2 different days |

**Supplementary Table 2.** Summary of mapping and filtering by CellRanger count. The sample IDs are the same as the ones used in the Seurat object. The number of cells is estimated by filtering out empty droplets by a computed threshold at the elbow of the barcode rank plot. The detailed summary outputs from CellRanger are available upon request.

| Sample | Estimated Number of Cells | Mean Reads per Cell | Median Genes per Cell | Fraction Reads in Cells | Confidently Mapped to the Transcriptome | Number of Reads |
| --- | --- | --- | --- | --- | --- | --- |
| **ML231-B2** | 7,494 | 33,224 | 1,031 | 50.4% | 43.7% | 248,979,225 |
| **ML232-B2** | 9,094 | 29,972 | 1,602 | 69.8% | 64.8% | 272,561,325 |
| **ML232-B1** | 9,087 | 29,822 | 1,667 | 70.2% | 63.4% | 270,990,370 |
| **ML232-A2** | 7,052 | 30,717 | 1,621 | 71.3% | 64.6% | 216,616,437 |
| **ML245-A1** | 11,001 | 26,380 | 1,609 | 75.0% | 55.1% | 290,211,004 |
| **ML245-D** | 21,648 | 22,588 | 1,621 | 73.3% | 59.1% | 488,974,966 |
| **ML263-C2** | 4,346 | 92,592 | 1,292 | 46.5% | 62.2% | 402,405,588 |
| **ML275-C1** | 8,535 | 28,833 | 1,278 | 56.2% | 60.1% | 246,088,402 |

**Supplementary Table 3.** Published pattern of expression of genes in *B.schlosseri*, with their ID in our dataset and publication of origin.

| Published gene name | Gene id and name in our dataset | Tissue/Cell type from budlet stage C1 | References |
| --- | --- | --- | --- |
| *Vasa* | *Boschl.Bs4.g5801-DDX4* | Germline | (32) |
| *Piwi* | *Boschl.Bs5.g6728-PIWIL1.1* | Germline | (30) |
| *Myh3* | *Boschl.Bs15.g19623-Myh6* | Body wall muscle | (23) |
| *Mrf* | *Boschl.Bs4.g5757-MYOD1* | Cells of the mantle (possibly muscle precursors) | (23) |
| *Ebf* | *Boschl.Bs10.g14018-EBF1* | Body wall muscle precursors and developing NS | (23) |
| *Myh2* | *Boschl.Bs8.g10861-MYH3.1* | Heart | (23) |
| *FoxF* | *Boschl.Bs7.g9951-FOXF1.1* | Epidermis + Dorsal branchial + Heart | (23) |
| *Tbx1/10* | *Boschl.Bs2.g1884-TBX1.1* | Mesenchymal cells+ branchial domain + the cerebral ganglion region + Forming heart | (23) |
| *Nk4* | *Boschl.Bs6.g8341-NKX2-3* | left branchial and left peribranchial, developing myocardium | (23) |
| *Gata-a* | *Boschl.Bs2.g2024-GATA5.1* | gut primordium, developing heart | (11) |
| *Tgf-β-f* | *Boschl.Bs8.g10511* | Follicular cells /Support cells | (32) |
| *Gata-b* | *Boschl.Bs13.g17419-GATA3.1* | peribranchial & posterior branchial | (11) |
| *FoxA1* | *Boschl.Bs6.g7874-FOXA1* | Gut rudiment (esp. oesophagus),<br>Endostyle rudiment, stigmata | (11) |
| *FoxA2* | *Boschl.Bs8.g11479-FOXA2* | Gut | (11) |
| *Zic-r.a* | *Boschl.Bs13.g17465* | developing CNS | (22) |
| *Pax3/7* | *Boschl.Bs8.g10830-PAX3.1* | developing and mature CNS | (22) |
| *Tbx2/3* | *Boschl.Bs9.g12221-TBX2.1* | Dorsal tube, Stomach, Branchial<br>flanking the endostyle | (11) |
| *SoxB1* | *Boschl.Bs11.g14812-SOX3.1* | Stigmata, Endostyle, Neural Gland,<br>Oesophagus, Germline, Heart, some<br>hemoblasts | (60) |

**Supplementary Table 4.**
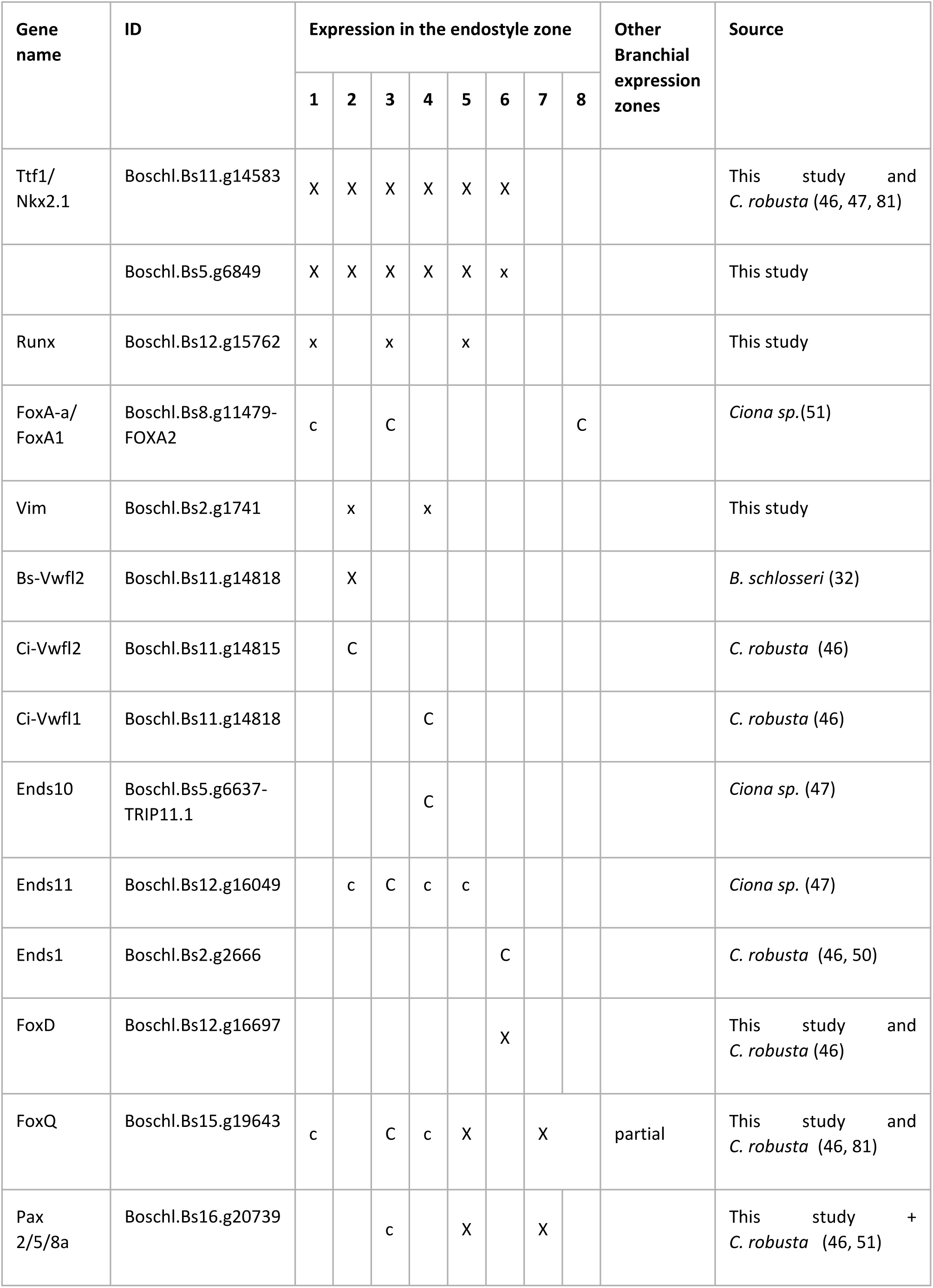
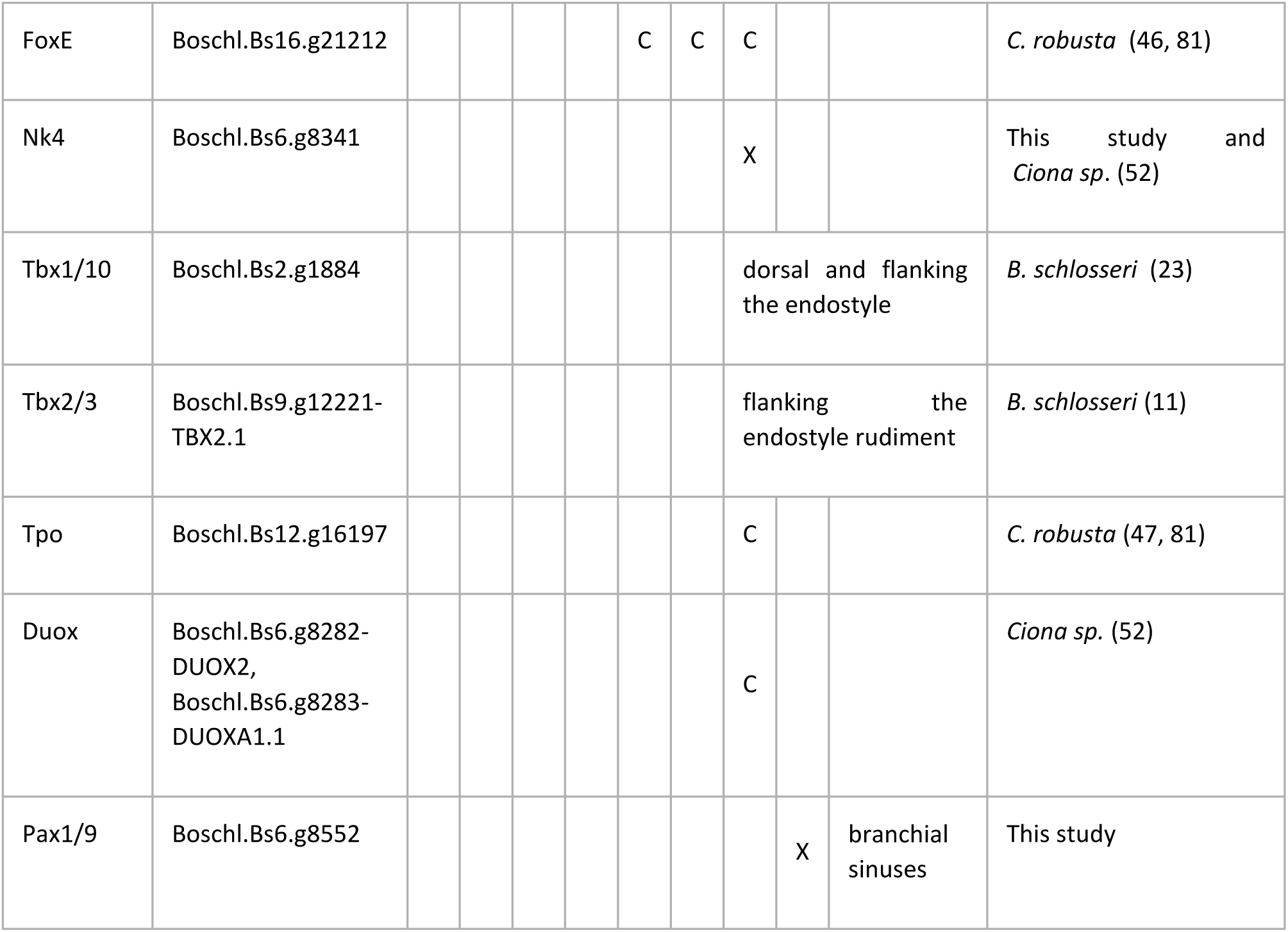
Published expression patterns in the endostyle zones and other branchial tissues. Genes used as reference for identifying the clusters in Fig. 3D. X/x expression pattern reported in *B. schlosseri*, C/c: expression reported in *Ciona* but not observed in this study. Uppercase: strong expression, lowercase: low/partial expression.

